# Granzyme B-based CAR T cells block metastasis by eliminating circulating tumor cells

**DOI:** 10.1101/2024.03.18.585442

**Authors:** Bin Sun, Jing Guo, Dong Yang, Qiancheng Hu, Haiyan Ma, Panwen Tian, Nan Liu, Longbao Lv, Lanzhen Yan, Hao Ding, Maoyong Fu, Hongfeng Gou, Dan Cao, Dan Liu, Nianyong Chen, Peng Shi, Weimin Li, Xudong Zhao

**Affiliations:** Division of Abdominal Tumor Multimodality Treatment and Laboratory of Animal Tumor Models, Cancer Center, State Key Laboratory of Respiratory Health and Multimorbidity, Frontiers Science Center for Disease-Related Molecular Network, West China Hospital, Sichuan University, Chengdu, Sichuan, 610041, China; Department of Pulmonary and Critical Care Medicine, Institute of Respiratory Health, State Key Laboratory of Respiratory Health and Multimorbidity, Frontiers Science Center for Disease-related Molecular Network, Precision Medicine Key Laboratory of Sichuan Province, West China Hospital, West China School of Medicine, Sichuan University, Chengdu, Sichuan, 610041, China; State Key Laboratory of Genetic Resources and Evolution, Kunming Institute of Zoology, Chinese Academy of Sciences, Kunming, Yunnan, 650223, China; Division of Abdominal Tumor Multimodality Treatment, Cancer Center, West China Hospital, Sichuan University, Chengdu, Sichuan, 610041, China; Division of Head & Neck Tumor Multimodality Treatment and Department of Radiation Oncology, Cancer Center, West China Hospital, Sichuan University, Chengdu, Sichuan, 610041, China; Department of Thoracic Surgery, Affiliated Hospital of North Sichuan Medical College, Nanchong, Sichuan, 637000, China

**Author notes:** Corresponding authors Xudong Zhao, No. 37 Guoxue Alley, Chengdu, Sichuan, 610041, China. These authors contributed equally (Bin Sun, Jing Guo, Dong Yang, Qiancheng Hu).

**Keywords:** mHSP70, Chimeric antigen receptor, Circulating tumor cells, Spontaneous metastasis models

## Abstract

Chimeric antigen receptor (CAR) T cells have limited efficacy against solid tumors due to the hostile microenvironment. Circulating tumor cells (CTCs) are essential to metastasis, which is the cause of most of cancer-related death. Here, we generated GrB-CAR T cells targeting membrane-bound HSP70 (mHSP70), a highly tumor-specific antigen detected in numerous cancers. GrB-CAR T cells exhibited potent cytotoxicity against a broad spectrum of cancer cell lines and stem-like cancer cells *in vitro* and effectively inhibited xenograft tumor growth *in vivo*. Importantly, GrB-CAR T cells markedly decreased the number of CTCs and, therefore, hindered cancer metastasis in spontaneous metastasis models with uncontrollable primary tumor growth, a scenario commonly encountered in clinical trials of CAR T therapies for solid tumors. Furthermore, despite the 100% homology between human and macaque HSP70 protein, the autotransplantation of macaque T cells expressing human GrB-CAR did not cause any obvious toxic effects. These results not only demonstrate GrB-CAR T cells as a safe and effective tactic with broad-spectrum anticancer activity, but also offer strong experimental evidence and proof-of-concept validation for CAR T cell-mediated metastasis inhibition by targeting CTCs.

## Introduction

Chimeric antigen receptor (CAR) T cells have presented a revolutionary opportunity to target cancer cells selectively and effectively to improve patient survival and have achieved considerable success in the treatment of hematologic malignancies, as demonstrated by the FDA approval of several CAR T-cell products targeting CD19 or BCMA[1]. Unfortunately, the lack of tumor-exclusive antigen targets and the extremely immunosuppressive and metabolically demanding tumor microenvironment limit the effectiveness of CAR T-cell therapies for solid tumors, which leads to insufficient tumor infiltration by CAR T cells as well as T-cell fatigue and dysfunction. Thus, there is an urgent need to develop new strategies to overcome these limitations in the context of solid tumors.

Metastasis is responsible for up to 90% of cancer-related fatalities, and circulating tumor cells (CTCs) are the main factor in the metastatic spread of solid tumors. CTCs are shed from the primary tumor *in situ*, intravasate and survive in the circulatory system, and enable the development of distal metastases[2]. A high CTC blood count has been linked to poor prognosis and an increased likelihood of metastasis in patients with solid tumors, such as breast[3], prostate[4], colorectal[5], lung[6], pancreatic[7] cancers, etc. Thus, targeting CTCs is a potential strategy to prevent or inhibit solid tumor metastasis. The immunosuppressive tumor microenvironment would not limit strategies targeting solid tumor-derived CTCs with CAR T cells; instead, they would mimic the successful targeting and killing of hematological tumor cells by CAR T cells and thus effectively block solid tumor metastasis by preventing CTCs from reaching distal sites. Nevertheless, intravenous tumor cell injections were unable to precisely simulate the shedding of cells from the original mass and replicate the the characteristics of CTCs in the earlier studies on the functions of CAR T cells in the prevention of cancer spread. In addition, in most of these studies, CAR T cells were injected several days after tumor cell injection so that the prevention of metastasis by CAR T cells was due to the elimination of micrometastases rather than CTCs[8-10]. In addition, the effects of primary tumor shrinkage on reducing the CTC number and suppressing metastasis need to be excluded in spontaneous metastasis models evaluating the roles of CAR T cells in CTCs and metastasis.

Heat-shock protein 70 (HSP70, Hsp72, or HSP70-1) encoded by the HSPA1A gene is expressed at low or undetectable levels in normal, unstressed cells[11]. However, it is highly induced by cellular stress to restore cellular homeostasis by expediting protein folding and the degradation of misfolded proteins as a chaperone protein. HSP70 is abundantly expressed in various types of human cancer cells to help cancer cells successfully respond to endogenous and exogenous stress, and HSP70 depletion in cancer cells induces cell death[12, 13]. Crucially, HSP70 is normally cytoplasmic in normal cells but can translocate to the cancer cells’ plasma membrane by binding to lipids unique to tumors, such as phosphatidylserine or globotriaosylceramide (Gb3)[14, 15], thus becoming accessible to CAR T cells. Consequently, the overexpression and specific cell surface localization of HSP70 in cancer cells could render membrane-bound HSP70 (mHSP70) an ideal tumor-specific target for CAR T-cell-based therapy, with a low risk of toxicity to normal cells. The metastatic tumors express higher mHSP70 than primary tumors, and mHSP70 was also found to be expressed on CTCs more frequently than EpCAM, which is an ideal CTCs target protein[16, 17].

To target mHSP70-expressing cancer cells, we generated CAR T cells based on natural ligand granzyme B (GrB-CAR T). The GrB-CAR T cells effectively killed cancer cells *in vitro* and inhibited xenograft tumor growth *in vivo*. Furthermore, they exhibited potent inhibition of mHSP70-expressing CTC-driven tumor metastasis in spontaneous metastasis models, even when the primary tumor mass could not be shrunk by CAR T cells, a common clinical situation encountered in CAR T therapy for solid tumors. GrB-CAR T cells did not cause any obvious side effects in mice or the nonhuman primate rhesus macaque, although human GrB has been reported to recognize mouse mHSP70 and human/rhesus macaque HSP70 share 100% homology. These results present a safe and efficient CAR T therapy targeting mHSP70 and the first well-designed experimental evidence of metastasis control by elimination of CTCs via CAR T cells.

## Results

### GrB-based CAR T cells efficiently target mHSP70-positive cancer cells

The expression and localization of mHSP70 were evaluated in multiple human cancer cell lines, including the pancreatic cancer cell lines PANC-1, AsPC-1, BxPC-3, and MIA PaCa-2, the liver cancer cell lines SMMC-7721, SK-Hep-1, and HepG2; and other cell lines, including small-cell lung cancer (SCLC) cell lines NCI-H1339 and NCI-H446, non-small cell lung cancer cell line NCI-H1299, prostate cancer cell line PC-3, glioblastoma cell line U-87 MG, colorectal cancer cell line HCT116, and breast cancer cell line MCF7. All the tested cancer cell lines expressed mHSP70 except for PANC-1 (**Fig. 1a** and **Supplementary Fig. 1**). However, the level of total HSP70 protein in PANC-1 cells was comparable to that in the other tested pancreatic cancer cell lines, as determined by Western blotting (**Fig. 1b**), suggesting that the translocation mechanism of HSP70 is independent of protein expression.

**Fig. 1.**
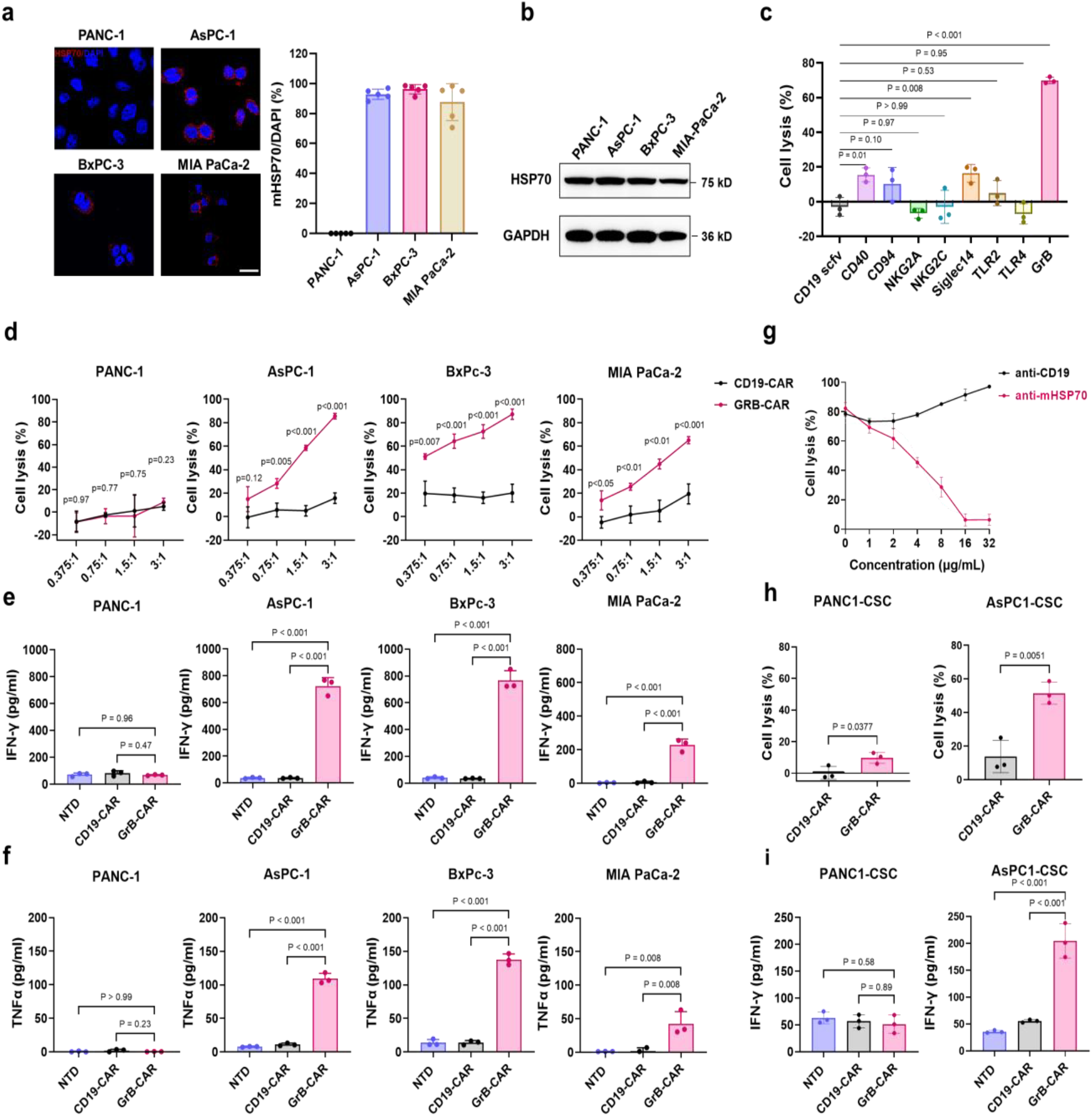
GrB-based CAR T cells efficiently target mHSP70-positive cancer cells. **a**. Representative image of mHSP70 expression in various cancer cell lines (left). Scale bar, 20 μm. The percentage of mHSP70-positive cells was quantified (right), n=5 independent experiments. **b**. Detection of HSP70 protein in human pancreatic cancer cells by western blotting. GAPDH was used as a loading control. **c**. Cytotoxicity of the indicated CAR T cells against HepG2 cells. **d**. Cytotoxicity assay of GrB-CAR T cells. CAR T cells were incubated with luciferase-expressing PANC-1, AsPC-1, BxPC-3, or MIA PaCa-2 cells for 16 hours at different effector to target (E: T) ratios. The in vitro cytotoxicity of the CAR T cells was determined by a bioluminescence assay. **e-f**. The concentrations of IFN-γ (e) and TNF-α (f) in the supernatant of the coculture system at an E:T ratio of 3:1, as measured by ELISA. **g**. Cytotoxicity assay of GrB-CAR T cells against AsPC-1 cells in the presence of either CD19-or mHSP70 antibody at an E:T ratio of 3:1. **h**. Cytotoxicity of GrB-CAR T cells against stem-like AsPC-1 and PANC-1 cells cultured at an E:T ratio of 3:1. **i**. Release of IFN-γ as measured by ELISA in the supernatants of stem-like AsPC-1 and PANC-1 cells treated with control or CAR T cells. Data in c-i are representative of three biological replicates from n=2 independent experiments. The results are presented as the means ±SDs.

To target mHSP70-positive cancer cells, we constructed lentiviral vectors containing a series of second-generation CARs comprising a CD8 signal peptide, mHSP70 ligands as the recognition domain, the CD8 hinge and transmembrane region, and the 4-1BB and CD3ζ signaling domains (**Supplementary Fig. 2a**) in which the mKate2 fluorescent protein was fused to the C-terminus of each CAR via a T2A sequence as a marker of T-cell transfection. A scFv CAR targeting CD19 (CD19-CAR) was used as a negative control. The lentiviral transduction efficiency in T cells ranged from 12% to 38%, as monitored by flow cytometric analysis of mKate2 expression (**Supplementary Fig. 2b**). The transduced T cells were tested for cytotoxicity against the mHSP70-positive HepG2 cell line, and the GrB-based CAR showed the highest cytotoxic efficiency (**Fig. 1c)**. Therefore, we chose GrB-CAR T cells for further study.

**Fig. 2.**
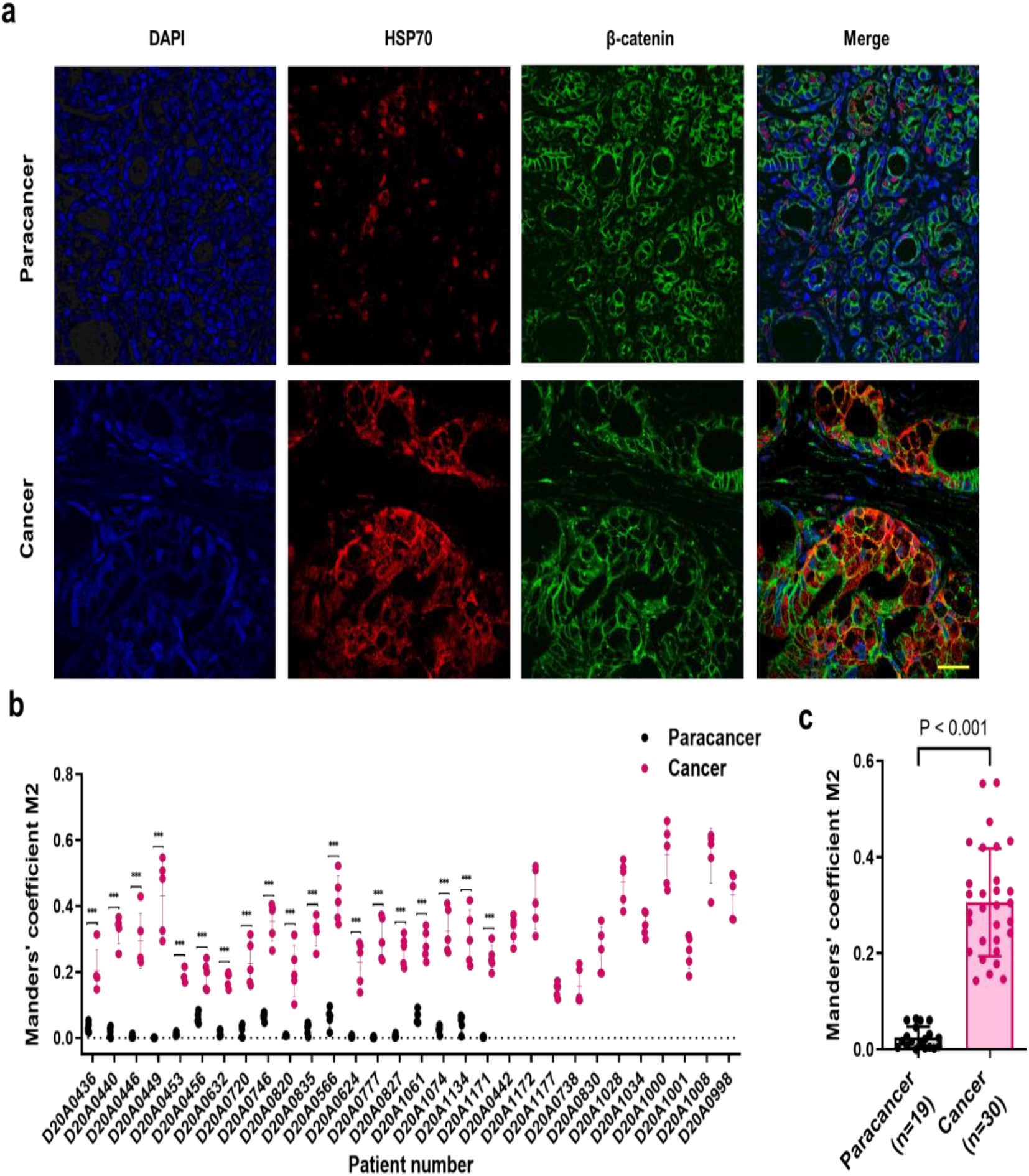
The expression of mHSP70 in human pancreatic tumor tissues. **a**. Representative immunofluorescence images of pancreatic tumor tissues stained with anti-HSP70 (red) and anti-β-catenin (green) antibodies; nuclei were counterstained with DAPI (blue). Colocalization of HSP70/β-catenin in tumor tissues is shown in the merged images. Scale bar, 25 μm. **b**. HSP70/β-catenin colocalization in tumor tissues was quantified based on Manders’ coefficient M2 in five images per sample. Nineteen pairs of paracancerous and pancreatic tumor tissues were included, along with individual informative pancreatic tumor samples. **c**. Statistical analysis of Manders’ coefficient M2 for the paracancerous tissues (n=19) and the pancreatic cancer tissues (n=30). The data are presented as the means ±SDs.

We assessed the cytotoxicity of GrB-CAR T cells against a variety of mHSP70-expressing cancer cells, especially pancreatic cancer cell lines. The function of GrB-CAR T cells was tested *in vitro* using a luciferase-based cytotoxicity assay. Pancreatic cancer cells were potently lysed by the CAR T cells in an mHSP70- and dose-dependent manner (**Fig. 1d**). Among the mHSP70-positive pancreatic cancer cell lines tested, namely, AsPC-1, BxPC-3, and MIA PaCa-2, the percentages of cells eliminated in response to GrB-CAR T cells were 87%, 85%, and 65%, respectively, at an effector: target (positive CAR T cell: target cell) ratio of 3:1. In contrast, GrB-CAR T cells did not affect mHSP70-negative PANC-1 cells (**Fig. 1d**). Consistently, the concentration of the cytokines IFN-γ and TNF-α in culture medium from the GrB-CAR T group was significantly increased compared to that in CD19-CAR T group (**Fig. 1e-f**), demonstrating the antigen-dependent activation of T cells. All the tested cell lines of other cancer types were also efficiently lysed by GrB-CAR T cells, accompanied by the release of cytokines (**Supplementary Fig. 3**). To further elucidate the specificity of GrB-CAR T cell cytotoxicity, an mHSP70 antibody made in our lab with reported Hsp70-derived TKD peptide as immunogen[18] was used to block the target of CAR T cells in a killing assay with ASPC-1 cells. The anti-mHSP70 antibody preincubation for one hour demonstrated, as Fig. 1g illustrates, a dose-dependent suppression of CAR-T cell cytotoxicity, with nearly total blocking shown at a concentration of 16 μg/ml. **(Fig. 1g)**. These findings demonstrate that GrB-CAR T cells selectively and effectively target a broad spectrum of cancer cells expressing mHSP70 *in vitro*.

**Fig. 3.**
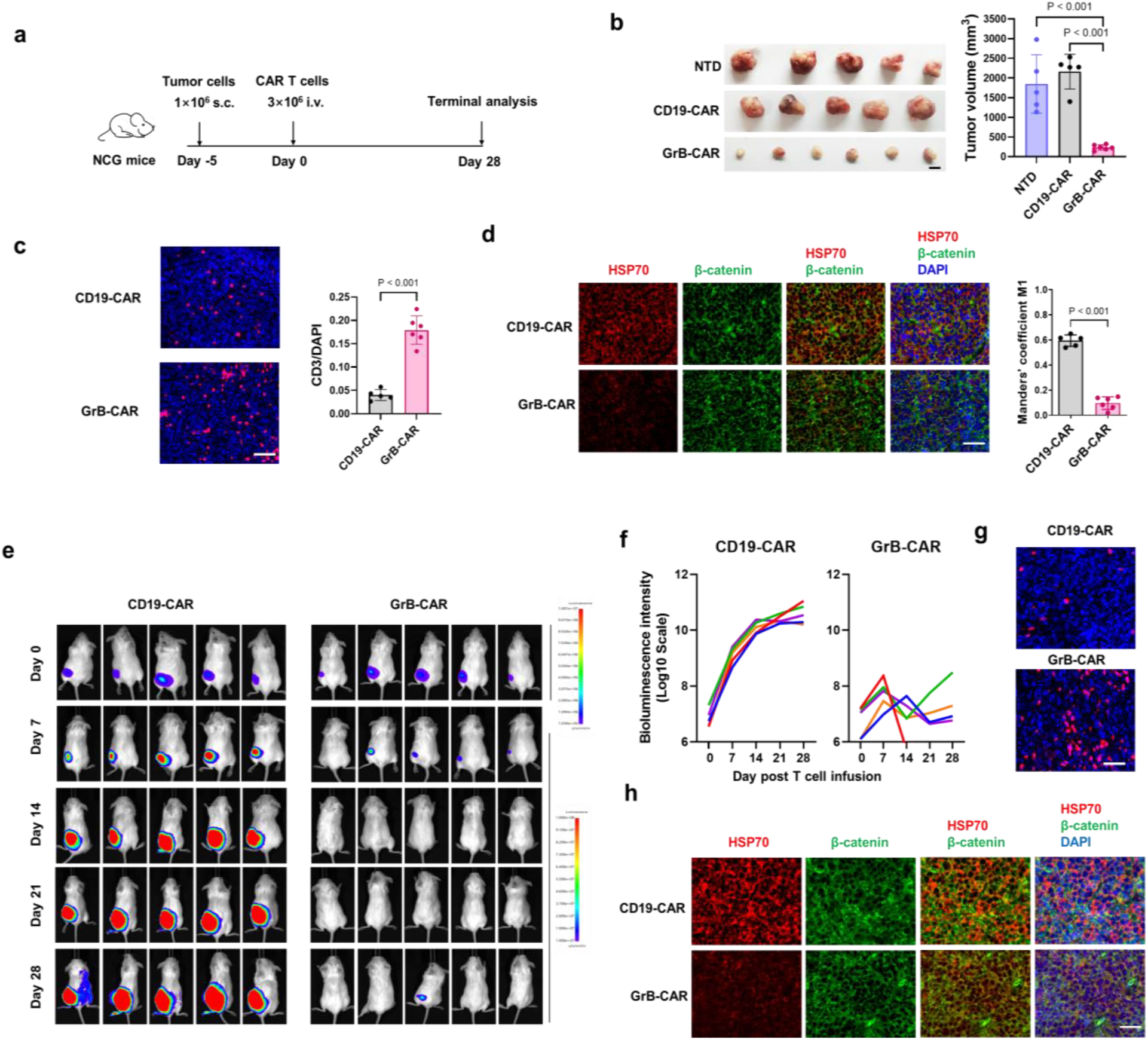
GrB-CAR T cells suppress tumor growth in xenograft mouse models. **a**. Schematic diagram of the establishment of the in vivo experimental model to determine the effect of GrB-CAR T cells on tumor growth. NCG mice were subcutaneously injected with 1×10^6^ AsPC1-luc cells, and 3×10^6^ CD19-CAR T cells or GrB-CAR T cells were then intravenously injected into the xenograft-bearing mice at the indicated time points. **b**. Effects of CAR T cells on the growth of xenograft tumors derived from the human pancreatic cancer cell line AsPC-1. Left, images of tumors collected from the NTD group (n=5), CD19-CAR T-cell group (n=5) and GrB-CAR T-cell group (n=6). Scale bar, 1 cm. Right, quantification of tumor volume as shown in the left panel. Each dot represents a tumor from an individual mouse. The data are presented as the means ±SDs and were analyzed by ANOVA with the Tukey’s correction for multiple comparisons. **c**. Representative images of immunofluorescence staining of CD3 in human pancreatic tumors treated with GrB-CAR T or CD19-CAR T cells (left). Scale bar, 50 μm. Quantification of CD3^+^ T cells per image field is shown as the mean ±SD (right). **d**. Representative images of immunofluorescence staining of mHSP70 and β-catenin in human pancreatic tumors treated with GrB-CAR T or CD19-CAR T cells (left). Scale bar, 50 μm. HSP70/β-catenin colocalization in tumor tissues was quantified based on Manders’ coefficient M1 in five images per sample (right). **e**. Bioluminescence imaging of mice bearing tumors derived from the human hepatocarcinoma cell line SMMC-7721 and treated with CD19-CAR T or GrB-CAR T cells. Each column shows one mouse over time. **f**. Curve of the bioluminescence intensity shown in (e). **g.** Representative images of CD3 immunofluorescence staining for the detection of CAR T cells in xenograft tumors. Mice bearing SMMC7721-derived xenograft tumors were sacrificed on day 28 after CAR T-cell treatment, and tumors were collected. Immunofluorescence staining of human CD3 was performed to detect infiltrated CAR T cells. Scale bar, 50 μm. **h**. Representative images of immunofluorescence staining of mHSP70 and β-catenin in xenograft tumors. Mice bearing xenograft tumors derived from the human hepatocarcinoma cell line SMMC-7721 were sacrificed on day 28 after CAR T-cell treatment, and tumor tissues were collected. Scale bar, 50 μm.

Growing data indicates that stem-like cancer cell is a useful target for inhibiting tumor development [19, 20]. To test the effects of GrB-CAR T cells on stem-like cancer cells, we generated stem-like tumorspheres from the human pancreatic cancer cell lines AsPC-1 and PANC-1 (named AsPC1-CSCs and PANC1-CSCs, respectively), which were confirmed by the induced expression of cancer stem cell (CSC) markers such as CD133, OCT4 and CXCR4 (**Supplementary Fig. 4a**). Consistent with the results in the parental cell lines, mHSP70 was detected on AsPC1-CSCs but not on PANC1-CSCs (**Supplementary Fig. 4b**). Correspondingly, GrB-CAR T cells exhibited potent cytotoxicity and induced robust release of IFN-γ from coculture with AsPC1-CSCs but not PANC1-CSCs (**Fig. 1h, i**), suggesting that CSCs express mHSP70 in a way similar to cancer cells and can be lysed by GrB-CAR T cells.

**Fig. 4.**
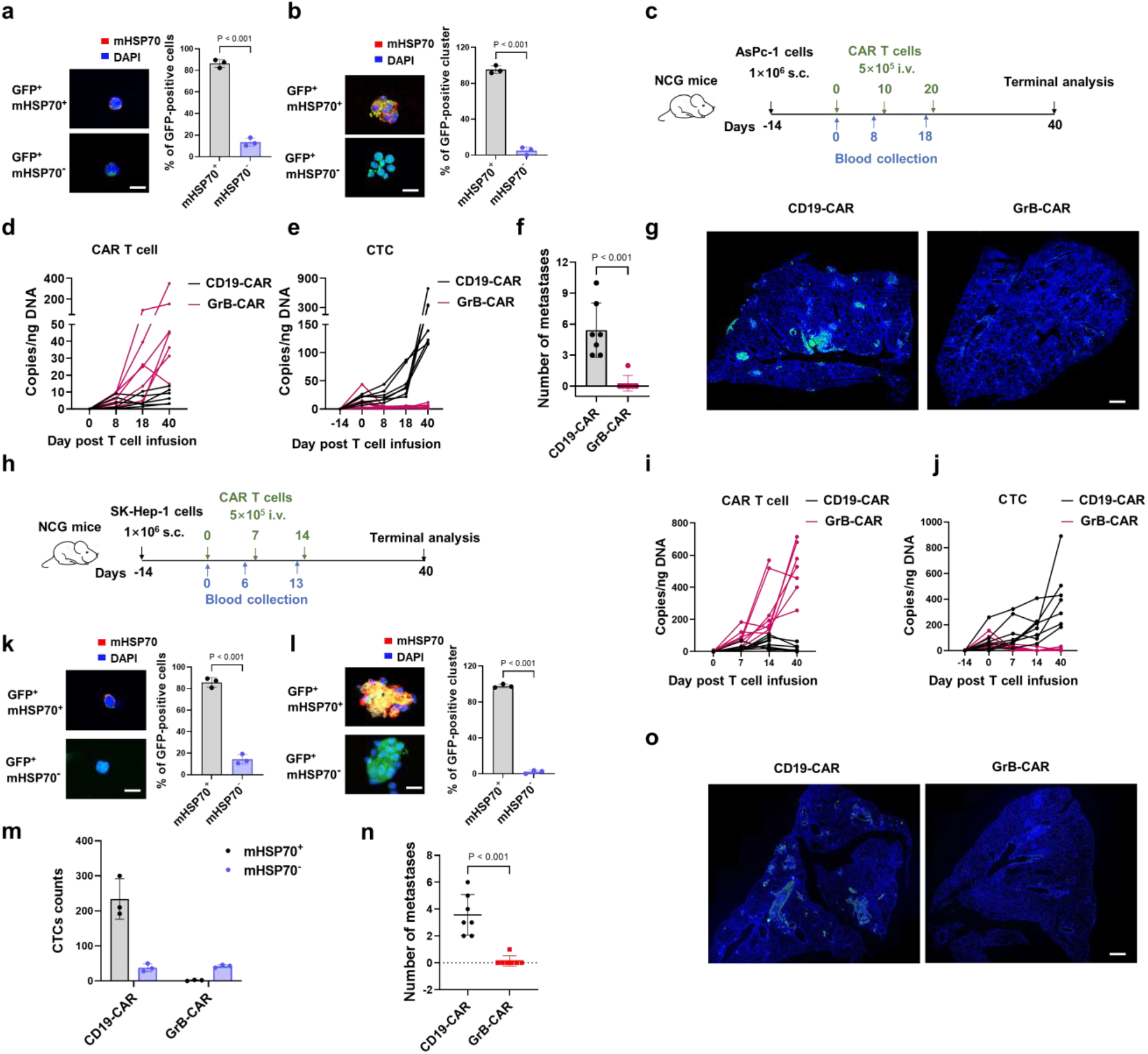
GrB-CAR T cells target CTCs to suppress tumor metastasis in vivo. **a-b**. NCG mice were subcutaneously injected with GFP-labeled AsPC1 cells. Fifty-two days later, CTCs were collected from the peripheral blood of the mice by GFP fluorescence for mHSP70 immunofluorescence staining. The expression of mHSP70 on total CTCs (**a**) and CTC clusters (**b**) was quantified. Each dot represents a single mouse, n=3. Scale bars for (a), 10 μm; Scale bars for (b), 20 μm. **c**. Schematic diagram of the establishment of the AsPC-1 xenograft model to determine the effects of GrB-CAR T cells on tumor metastasis. NCG mice were subcutaneously injected with 1×10^6^ AsPC1-luc cells. Then, 5×10^5^ CD19-CAR T cells or GrB-CAR T cells were intravenously injected into the xenograft-bearing mice at the indicated time points. **d-e**. Quantification of DNA copies of CAR T cells (**d**) and CTCs (**e**) in peripheral blood collected from tumor-bearing mice, n=7 per group. **f**. Number of metastases on the lung surface in NCG mice treated with CD19-or GrB-CAR T cells. Each dot represents the metastatic nodules in an individual mouse, n=7 per group. **g**. Metastasis of human tumor cells in mouse lung tissue upon CAR T-cell challenge as indicated. Representative images of immunofluorescence staining with a human-specific anti-COXIV antibody (green) and DAPI (blue) in the lungs. Scale bars, 500 μm. **h**. Schematic diagram of the establishment of the SK-Hep-1 xenograft model to determine the effects of GrB-CAR T cells on tumor metastasis. NCG mice were subcutaneously injected with 1×10^6^ SK-Hep1-luc cells. Then, 5×10^5^ CD19-CAR T cells or GrB-CAR T cells were intravenously injected into the xenograft-bearing mice at the indicated time points. **i**-**j**. Detection of DNA copies of CAR T cells (**i**) and CTCs (**j**) in peripheral blood collected from tumor-bearing mice. n=8 per group. **k**-**m**. CTCs were collected from mouse peripheral blood for mHSP70 immunofluorescence staining on day 40 following treatment with CD19-CAR or GrB-CAR T cells. The expression of mHSP70 on total CTCs (**k**) and CTC clusters (**l**) isolated from CD19-CAR T cell-treated mice was quantified. Statistics on the number of total mHSP70^+^ and mHSP70^-^ CTCs in the mice treated by CD19-CAR or GrB-CAR T cells (**m**). Each dot represents a single mouse, n=3. Scale bars for (k), 10 μm; Scale bars for (l), 20 μm. **n**. Metastases in the lungs. Number of metastases on the surface of lungs harvested from mice on day 40 after treatment with CD19-CAR (n=7) or GrB-CAR (n=8) T cells. Each dot represents the metastatic nodules in an individual mouse. **o**. Metastatic nodules derived from human tumor cells in mouse lung tissue upon CAR T-cell challenge as indicated. Representative images of immunofluorescence staining with a human-specific anti-COXIV antibody (green) and DAPI (blue) in the lungs. Scale bars, 500 μm. The data are shown as the means ±SDs in a,b,f,k-n.

While it has been noted that mHSP70 is expressed in most malignancies but not in normal tissues[21], specific cancer types still need more investigation. To characterize the membrane localization of HSP70 in pancreatic cancer, double immunohistochemical staining for HSP70 and the membrane marker β-catenin was performed with a TSA fluorescence system on a cancer tissue chip (HPanA060CS02, Shanghai Outdo Biotech Co., Ltd., China) containing 23 pairs of cancer and paracancerous tissues and 14 additional cancer samples. Colocalization of HSP70 with β-catenin on the membrane (membrane-bound HSP70) was widely observed in cancer tissues (**Fig. 2a** and **Supplementary Fig. 5**). However, colocalization was rare in paracancerous tissues, although some intracellular HSP70 staining was observed **(Fig. 2a)**. After excluding samples with poor tissue or image quality, the remaining 19 tissue pairs and an additional 11 cancer samples were used for quantitative analysis of the colocalization of HSP70 and β-catenin. mHSP70 expression was significantly higher in each tumor sample than in the paired paracancerous tissue sample and in the overall comparison containing paired and unpaired samples (**Fig. 2b, c**). The expression of mHSP70 in a variety of normal tissues was also examined utilizing a tissue chip (HorgN120PT01, Shanghai Outdo Biotech Co., Ltd., China). Although some staining was visible in a few samples, including the esophagus, urinary bladder, colon, lung, testis, epidermis, epididymis, and prostate samples, the signal was predominantly intracellular and did not overlap with membrane β-catenin (**Supplementary Fig. 6**).

**Fig. 5.**
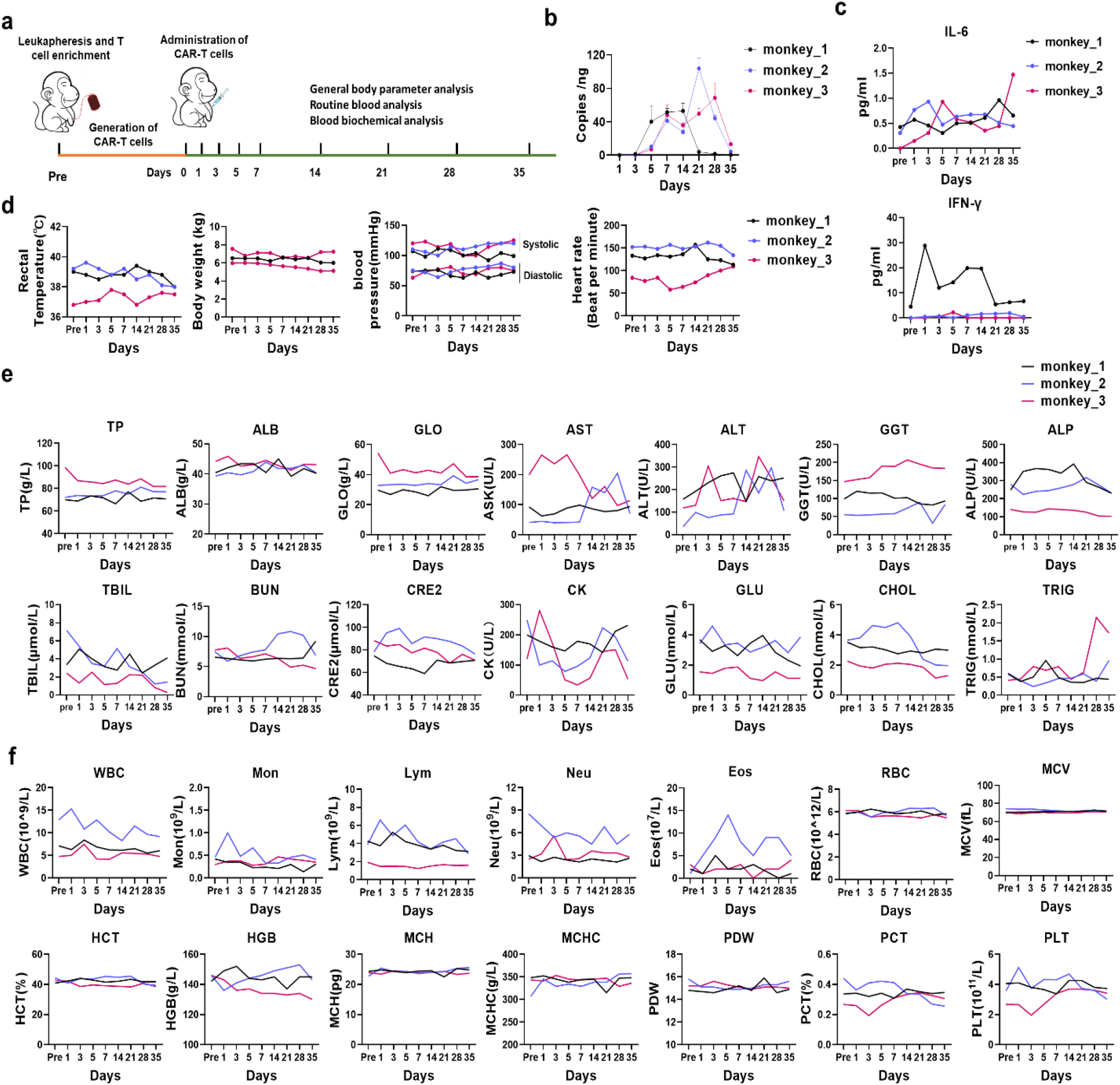
Safety evaluation of GrB-CAR T cells in nonhuman primates. **a**. Schematic diagram of the experimental design for safety evaluation. **b**. DNA copies of GrB-CAR T cells in the peripheral blood of macaques were quantified by real-time PCR. **c**. Quantitation of plasma concentrations of IL-6 and IFN-γ in blood collected from nonhuman primates using MesoScale Discovery V-Plex assay kits. **d**. The body weight, rectal temperature, blood pressure, and heart rate of three macaques were measured at the indicated times after T-cell infusion. **e**. Blood biochemical analysis was performed to measure total protein (TP), albumin (ALB), globulin (GLO), aspartate aminotransferase (AST), alanine aminotransferase (ALT), glutamyl transpeptidase γ-glutamyl transpeptidase (GGT), alkaline phosphatase (ALP), total bilirubin (TBIL), blood urea nitrogen (BUN), creatinine (CRE2), creatine kinase (CK), glucose (GLU), cholesterol (CHOL), and triglyceride (TRIG) at the indicated times after CAR T-cell transplantation. **f**. Hematological analysis was performed to determine the numbers of white blood cells (WBC), monocytes (Mon), lymphocytes (Lym), neutrophils (Neu), eosinophils (Eos), red blood cells (RBC) and mean corpuscular volume (MCV), hematocrit (HCT), hemoglobin (HGB), mean corpuscular hemoglobin (MCH), mean corpuscular hemoglobin concentration (MCHC), platelet distribution width (PDW), plateletcrit (PCT), and platelet count (PLT) at the indicated times after CAR T-cell transplantation.

### GrB-CAR T cells suppress tumor growth in xenograft mouse models

To investigate the antitumor effect of GrB-CAR T cells *in vivo*, 1×10^6^ AsPC-1 human pancreatic cancer cells were subcutaneously transplanted into immunodeficient NCG mice. Five d post tumor engraftment, the mice were divided into three groups with equal tumor sizes based on *in vivo* imaging of luciferase activity and infused intravenously with 3×10^6^ nontransduced (NTD) T cells, CD19-CAR T cells, or GrB-CAR T cells (**Fig. 3a**). The mice were sacrificed 28 d after treatment. The xenografts were harvested, and the volume of the tumors was analyzed. Compared with NTD T cells and CD19-CAR T cells, GrB-CAR T cells markedly inhibited tumor growth (P< 0.001, **Fig. 3b**). The tumors were sectioned and stained for CD3. The results showed significantly increased T-cell infiltration in residual tumors collected from GrB-CAR T-cell-treated mice (P< 0.001, **Fig. 3c**). To investigate the mHSP70 status of the tumors *in vivo*, double staining was performed for HSP70 and β-catenin. Opposite of T cell infiltration, very weak HSP70 expression was observed in the residual tumors compared to the tumors in CD19-CAR T-cell-treated mice (**Fig. 3d**), indicating that tumor immune escape was due to the loss of mHSP70 expression rather than GrB-CAR deficiency. Together, these results demonstrate the efficiency of GrB-CAR T cells in targeting and eliminating pancreatic cancer cells *in vivo*.

To evaluate the effects of GrB-CAR T cells in other xenograft models, NCG mice were subcutaneously inoculated with SMMC-7721 human hepatocarcinoma cells expressing luciferase. The luciferase signal increased with time, indicating that the tumors in mice treated with CD19-CAR T cells developed quickly. Conversely, the luciferase signal in GrB-CAR T-cell-treated mice was almost completely attenuated on d 7 after CAR T-cell infusion and was detectable in only one mouse on d 28 (**Fig. 3e, f**). The only residual tumor in the group treated with GrB-CAR T cells was analyzed. Similar to the results in AsPC-1 xenografts, numerous T cells were detected in the residual tumors in mice treated with GrB-CAR T cells, whereas few T cells were observed in CD19-CAR T-cell-treated tumors (**Fig. 3g**). The lack of mHSP70-expressing cancer cells in the residual tumors in mice treated with GrB-CAR T cells suggested that GrB-CAR T cells efficiently killed mHSP70-positive cancer cells and therefore suppressed liver tumor growth *in vivo* (**Fig. 3h**).

### GrB-CAR T cells target CTCs to suppress tumor metastasis in spontaneous metastasis models

Previous investigations on the effects of CAR-T on metastasis have predominantly relied on experimental metastasis models, which did not recapitulate the CTCs shed from the primary tumor. The universal expression of mHSP70 on CTCs [17, 22] suggests that GrB-CAR T cells may eliminate CTCs. To evaluate the ability of GrB-CAR T cells to eliminate CTCs and inhibit metastasis in spontaneous metastasis models, we first assessed whether CTCs maintain mHSP70 expression in a subcutaneous xenograft experiment with GFP-expressing AsPC-1 cells in NCG mice. Consistent with AsPC-1 cells *in vitro* culture, most GFP-positive tumor cells among mouse peripheral blood mononuclear cells (PBMCs) were mHSP70-positive. (**Fig. 4a**). CTC clusters are thought to be much more metastatic than single CTCs[23]. Almost all GFP-positive cell clusters expressed mHSP70 (**Fig. 4b**). We also evaluated mHSP70 expression in the CTC-enriched population isolated from PBMCs of patients with pancreatic cancer. CTC-enriched cells were stained with a mixture of an anti-mHSP70 antibody and an anti-CD45 antibody and were then incubated with a human chromosome 8 (CEP8) FISH probe. The mHSP70-positive CTCs among CD45-negative cells were identified as multiple puncta of CEP8 signals (**Supplementary Fig. 7**). These results indicate that CTCs can theoretically be targeted by GrB-CAR T cells to block tumor metastasis *in vivo*.

In order to investigate this potential, we created a model of AsPC-1-derived spontaneous metastasis and carefully calibrated the quantity and time of GrB-CAR T-cell infusion to prevent CAR T cell-mediated *in situ* tumor suppression, which could potentially lower the creation of CTCs (**Supplementary Fig. 8a**). On d 14, we administered CAR T cells by a single injection in gradient doses of 1×10^6^, 2×10^6^ or 3×10^6^ cells per mouse into NCG mice bearing AsPC-1 xenografts. There was no significant difference in the growth of primary subcutaneous tumors (**Supplementary Fig. 8b**). Genomic DNA was purified from peripheral blood, and the DNA copy number of CTCs was measured by quantitative PCR. The clearance of CTCs by GrB-CAR T cells was dose-dependent, and all doses of CAR T cells significantly reduced the number of CTCs (**Supplementary Fig. 8c**). According to existing research, AsPC-1 xenografts are most likely to metastasize to the lungs[24]. Therefore, the metastatic tumor cells in lung tissue were further analyzed by immunofluorescence staining with a human COXIV-specific antibody thirty d after CAR T-cell treatment. Consistent with the results of the DNA copy number of CTCs in blood, very few tumor nodules were detected in the lungs after 1×10^6^ GrB-CAR T-cell treatment. Notably, no metastatic nodules were detected in the lung tissues of mice treated with higher doses of CAR T cells (**Supplementary Fig. 8d**). Multiple doses of a lower number of CAR T cells probably decrease the side effects of CAR T treatment. Next, we further optimized the number of GrB-CAR T cells to explore whether multiple treatments at lower doses can inhibit tumor metastasis (**Fig. 4c**). Three treatments with 5×10^5^ GrB-CAR T cells did not affect primary tumor growth under the current experimental conditions (**Supplementary Fig. 9**). CD19-CAR T cells were maintained at very low levels, while GrB-CAR T cells continued to expand as detected in blood samples (**Fig. 4d**). Conversely, the number of DNA copy number of CTCs in the CD19-CAR T group increased over time, whereas that in the GrB-CAR T group declined on d 8 and remained almost undetectable up to 40 d after GRB-CAR T-cell treatment (**Fig. 4e**). Forty d after CAR T-cell treatment, all animals were dissected and subjected to gross examination of metastatic tumors in major organs, including the liver, lung, spleen, pancreas, kidney, and small intestine. Metastatic tumors were found only in the lung. All the animals in the CD19-CAR T group had metastatic tumors, consistent with the increased DNA copy number of CTCs over time. Conversely, out of nine mice treated with GrB-CAR T cells, only two tiny metastatic tumors were discovered in one mouse; this finding was consistent with the extremely low DNA copy number of CTCs (**Fig. 4f**). The metastatic tumor cells in lung tissue were also analyzed using a human COXIV-specific antibody. The number and size of metastatic nodules in the lungs were obviously lower in the GrB-CAR T group than in the CD19-CAR T group (**Fig. 4g**). Furthermore, the expression of HSP70 was barely detectable in the distant metastases in the GrB-CAR T group, indicating that these nodules were derived from HSP70-negative cells (**Supplementary Fig. 10**).

We further investigated the potential of GrB-CAR T cells to prevent metastasis in an additional spontaneous metastasis model using luciferase-GFP-labeled SK-Hep-1 liver cancer cells (**Fig. 4h**). Similar to the results in the AsPC-1 model, the growth of xenograft tumors was not significantly affected by GrB-CAR T cells (**Supplementary Fig. 11**). However, the DNA copy number of CTCs in the blood was significantly reduced, and meanwhile the DNA copy number was significantly increased in GrB-CAR T cell treated group compared to those in the CD19-CAR group (**Fig. 4i, j**). The vast majority of single CTCs and CTC clusters in PBMCs isolated from the mice treated with CD19-CAR T cells expressed mHSP70 (**Fig. 4k, l**), as the expression of mHSP70 in SK-Hep-1 cells cultured *in vitro*. However, mHSP70-positive tumor cells were almost completely absent, and only a few mHSP70-negative tumor cells were detected in GrB-CAR T-cell-treated mice (**Fig. 4m**). GrB-CAR T cell-treated mice had less apparent metastatic nodules on their lung surfaces, which was consistent with a decrease in CTCs in their blood (**Fig. 4n**). In addition, the GrB-CAR T-cell-treated mice had much fewer and much smaller metastatic nodules in the lungs than the CD19-CAR T-cell-treated mice as detected by immunofluorescence staining with human COXVI antibody (**Fig. 4o**). Similar to metastases derived from AsPC-1 cells, only very few HSP70-positive cells were observed, indicating that these metastases originated from HSP70-negative cells (**Supplementary Fig.12**).

### GrB-CAR T cells do not induce severe side effects in mice and nonhuman primates

No obvious side effects were observed in the xenograft mice in Figure 3, although human GrB was previously reported to recognize mouse mHSP70[25, 26]. To further examine the acute distribution of CAR T cells and their potential toxicity, we infused GrB-CAR or CD19-CAR T cells into NSG mice bearing xenograft tumors derived from AsPC-1 cells and collected tissues from major organs, such as the lung, liver, spleen, brain, pancreas, heart, and kidney, on d 5 after injection. GrB-CAR T cells accumulated abundantly in the xenograft tumors, whereas few CD19-CAR T cells were detected in the tumors (**Supplementary Fig. 13**). Additionally, a small number of CAR-engineered human T cells were found in the spleen and, to a lesser extent, in the lung and liver, but there were no appreciable differences between the GrB-CAR T-cell and CD19-CAR T-cell groups. No positive staining was found in the brain, pancreas, heart, or kidney (**Supplementary Fig. 13**). Furthermore, we did not find any obvious pathological lesions in any organs (**Supplementary Fig. 14**). These data demonstrate the specific infiltration of GrB-CAR T cells into tumors and the lack of obvious specific infiltration or tissue damage in normal organs.

Next, we assessed the safety of these cells in nonhuman primates. A schematic diagram of the experiment is shown in **Fig. 5a**. *Macaca mulatta* HSP70 protein (XP_014991489.2) shares 100% homology with human HSP70 protein (NP_005336), suggesting that a human GrB-based CAR could fully recognize mHSP70 in *M. mulatta*. To evaluate whether GrB-CAR T cells attack normal tissues in nonhuman primates, T cells were isolated from three adult *M. mulatta* macaques and stimulated with nonhuman primate-specific anti-CD2/CD3/CD28 Dynabeads prior to lentivirus-mediated GrB-CAR transduction. As monitored by mKate2 expression via flow cytometric analysis, the lentiviral transduction efficiency in T cells isolated from the three monkeys was 11%, 21.8%, and 26% (**Supplementary Fig. 15**). The macaques were administered autologous T cells armed with GrB-CAR. DNA was isolated from PBMCs collected at different time points and analyzed by quantitative PCR to quantify CAR T-cell expansion. The results showed that CAR T cells underwent moderate expansion *in vivo* (**Fig. 5b**). The concentrations of cytokines such as TNF-α, IFN-γ, IL-6, and IL-2 were measured in the plasma of the three macaques at various time points as an indication of CAR T cytotoxicity against normal cells; only IFN-γ and IL-6 were detectable in these three animals. Cytokine concentrations remained low after treatment with GrB-CAR T cells, although some variation in the IFN-γ concentration was observed in one macaque (**Fig. 5c**). The rectal temperature, body weight, blood pressure, and heart rate were monitored for 35 d and no significant changes were found (**Fig. 5d**). To further study the effects of GrB-CAR T cells on normal tissues, serum biochemical tests were performed to assess liver, heart, and kidney function, as well as metabolism-related serum parameters. These parameters including total protein (TP), albumin (ALB), globulin (GLO), aspartate aminotransferase (AST), alanine aminotransferase (ALT), glutamyl transpeptidase γ-glutamyl transpeptidase (GGT), alkaline phosphatase (ALP), and total bilirubin (TBIL) for liver function; blood urea nitrogen (BUN) and creatinine (CRE2) for kidney function; creatine kinase (CK) for cardiovascular function; and glucose (GLU), cholesterol (CHOL), and triglyceride (TRIG) for metabolic function were all within normal ranges (**Fig. 5e**). No abnormal changes in hematological parameters, such as the numbers of white blood cells (WBCs), monocytes (Mons), lymphocytes (Lyms), neutrophils (Neus), eosinophils (Eos), and red blood cells (RBCs) and mean corpuscular volume (MCV), hematocrit (HCT), hemoglobin (HGB), mean corpuscular hemoglobin (MCH), mean corpuscular hemoglobin concentration (MCHC), platelet distribution width (PDW), platelet hematocrit (PCT), and platelet count (PLT), were observed after CAR T-cell treatment (**Fig. 5f**). These findings indicate that in nonhuman primates, GrB-CAR T cells rarely attack normal cells *in vivo* and produce clear on-target/off-tumor effects.

## Discussion

The strict specificity of mHSP70 expression in cancer cells, including CTCs, makes it an ideal target for CAR T cell therapy. Although CAR T cells carrying an anti-mHSP70 single-chain variable fragment (scFv) kill colorectal cancer cells *in vitro*^[27]^, more sophisticated studies on the development of CAR T cells targeting mHSP70 are needed. Here, we demonstrated that GrB-CAR T cells exhibit cytotoxicity specifically in multiple cancer cell lines *in vitro* and significantly inhibit the growth of primary xenografts in vivo. More importantly, GrB-CAR T cells effectively decreased the number of CTCs and consequently inhibited cancer metastasis *in vivo* in spontaneous metastasis models, even in the presence of uncontrollable primary tumor growth, a common situation encountered in the clinical application of CAR T cells in solid tumor treatment. Moreover, GrB-CAR T cells did not elicit severe side effects in mice or nonhuman primates. These results show that GrB-CAR T cells are efficient at targeting a variety of mHSP70-positive cancer cells while preserving a good safety profile and that using CAR T cells to target CTCs is a viable tactic for treating solid tumors.

The utilization of a protein as a target for CAR T cell therapy requires cell surface expression. mHSP70 specifically express in majority of cancers and no expression was found in normal tissues. However, we found that mHSP70 expression on the cell surface did not necessarily correlate with the level of HSP70 expression in cancer cells. Total HSP70 protein levels in PANC-1 cells were comparable to those in other pancreatic cancer cell lines, but mHSP70 was not detected in PANC-1 cells. Consistently, GrB-CAR T cells were unable to lyse PANC-1 cells, highlighting the specificity of GrB-CAR T cells for cancer cells with mHSP70 expression. Therefore, it is crucial to identify mHSP70 expression rather than the entire HSP70 protein in order to direct the use of GrB-CAR T cells for cancer therapy.

Numerous studies have shown that CTCs are essential for metastasis, particularly when they are clustered and exhibit stemness and a hybrid epithelial-mesenchymal phenotype. The number and subclassification of CTCs have been well established as markers for prognosis and decision-making for therapies[28]. In addition, some strategies for blocking metastasis by targeting the molecular and cellular features of CTCs and CTC clusters have been tested in preclinical experiments[29, 30]. Na-K+ ATPase inhibitors were found to destroy CTC clusters to block metastasis[31], and this finding led to a clinical trial in breast cancer patients with progressive disease (NCT03928210) with the FDA-approved cardiac glycoside digoxin. However, CTC targeting remains challenging due to increased drug resistance, escape from immune surveillance, and constant release from the primary/metastatic tumor masses. CAR-engineered immune cells, especially long-lived T cells, offer a promising approach for enhancing immunosurveillance against CTCs. Although many studies have reported the blockade of metastasis by CAR T cells, these studies either employed experimental metastasis models that cannot accurately reproduce spontaneous metastasis in patients or only demonstrated the ability of CAR T cells to eliminate micrometastases by injection of these cells several days after tumor cell injection[8-10, 32]. As a result, there is still a lack of strong data supporting the effect of CAR T cells on CTCs. Previous studies identified mHSP70 as a stable cell surface marker of both epithelial and mesenchymal CTCs in several cancers[17, 22]. In this study, we established spontaneous metastasis models that were optimized to exclude the influence of primary tumor size on the abundance of CTCs during CAR T treatment. Our results strongly suggest that low doses of GrB-CAR T cells can effectively eliminate mHSP70-expressing CTCs shedding from primary tumors, thereby inhibiting tumor metastasis. Considering the widespread expression of mHSP70 in cancer samples, GrB-CAR T cells are probably a good choice for blocking metastasis by eliminating CTCs in many cancers. Although the therapeutic effect of CAR T cells on primary or metastatic tumor masses is currently limited, targeting CTCs with CAR T cells may avoid the difficulties presented by the immunosuppressive tumor microenvironment. This proof-of-concept study also provides a basis to extend the use of CAR T cells to prevent metastasis.

The most effective cancer treatments are still conventional chemotherapies and radiation treatments, which effectively reduce tumor bulk but also raise HSP70 and mHSP70 expression, which increases metastasis and leads to therapeutic resistance[33-35]. This provides an excellent rationale for combining traditional chemotherapies/radiotherapies with mHSP70-directed CAR T cell therapy. Together with the robust shrinkage of the tumor mass and alteration of the tumor microenvironment achieved by conventional chemotherapies/radiotherapies, the combination treatment with mHSP70-targeted CAR T cells is anticipated to yield promising outcomes of cancer treatment. The expression of mHSP70 on cancer cells can trigger an endogenous immune response, leading to the targeting of these cells by NK cells[36]. However, excessive expression of extracellular HSP70 can induce immunological tolerance and promote tumor progression[37-39]. Consequently, GrB-CAR T cells may offer a cutting-edge strategy to stop mHSP70-positive cancer cells from escaping the immune system.

The safety of CAR T cell therapy is a general concern in treating solid tumors. As far as we are aware, granzyme B has only one known receptor mHSP70[40], indicating that GrB-CAR T cells’ exclusive target is mHSP70. GrB itself can enter and induce kill cancer cells in either perforin-dependent or -independent manner^[25, 41]^ and can also be used to distribute functionalized superparamagnetic iron oxide nanoparticles in xenograft models with precision[26] while there were no obvious side effects observed in these xenograft mouse models. In this study, GrB in CAR molecule functions as the recognition domain for mHSP70 and cannot enter the cancer cells. Although human GrB exhibits activity on mouse cells[25, 26], GrB-CAR T-cell treatment did not result in any obvious toxic side effects in xenograft mouse models. Furthermore, macaques, being the nonhuman primates with high gene homology, similarity in the immune system and drug metabolism to humans[42, 43], serve as reliable models for assessing the safety of CAR T-cell therapy[44]. Autologous transplantation of *M. mulatta* T cells armed with GrB-CAR did not induce any serious side effects, although *M. mulatta* HSP70 shares 100% homology with human HSP70. Collectively, these data strongly suggest that GrB-CAR T cells are unlikely to cause serious or lethal on-target off-tumor effects in human patients. In addition, our experimental results demonstrated that CTCs might be eliminated with a comparatively smaller dose of GrB-CAR T cells, thus further lowering the possibility of adverse consequences.

GrB-CAR T cells may be a promising strategy advancing to clinical trials for treating multiple cancers. Strikingly, the elimination of CTC and blocking of metastasis by GrB-CAR-T cells may imply the earlier application of CAR-T in clinical trials to prevent possible metastasis. Our findings support additional investigations into CTC-targeting CAR T therapies for solid tumors to prevent metastasis.

## Materials and methods

### Cell culture

The human cancer cell lines U87-MG and HCT116 were purchased from the National Infrastructure of Cell Line Resource (Kunming, China). The MIA PaCa-2 and NCI-H1299 cell lines were purchased from the National Infrastructure of Cell Line Resource (Shanghai, China). The SK-Hep-1 cell line was purchased from Sichuan BIO Biotechnology Co., Ltd. (Chengdu, China). The NCI-H1339 cell line was purchased from Ningbo Mingzhou Biological Technology Co., Ltd. (Ningbo, China). The NCI-H466 cell line was a kind gift from Prof. Guangbiao Zhou (Cancer Hospital Chinese Academy of Medical Sciences). HepG2, MCF7, PC-3, SMMC-7721, AsPC-1, PANC-1, BxPC-3, and HEK-293T cells were maintained in our laboratory. PANC-1, MIA PaCa-2, HCT116, MCF7, HepG2, U87-MG, SK-Hep-1, PC-3, and HEK-293T cells were cultured in Dulbecco’s modified Eagle’s medium (DMEM; Life Technologies, USA), while AsPC-1, BxPC-3, NCI-H446, NCI-H1339, NCI-H1299, and SMMC-7721 cells were cultured in RPMI 1640 medium (Life Technologies) supplemented with 10% fetal bovine serum (FBS; Life Technologies) and 1× penicillin-streptomycin (Life Technologies). Luciferase-labeled cell lines were generated through infection with pTomo-CMV-luciferase-IRES-puro lentivirus (MOI≈10) and subsequently selected with puromycin (Life Technologies, 1 µg/ml) for 2 weeks. AsPC1-luciferase (AsPC1-luc) and PANC1-luc cells were cultured in serum-free stem cell medium consisting of DMEM/F12 (Life Technologies), EGF (20 ng/ml, Life Technologies), bFGF (Life Technologies, 20 ng/ml), and B27 (Life Technologies) to obtain cancer stem cell-like cells, namely, AsPC1-CSCs and PANC1-CSCs. All cells were cultured at 37 °C in a humidified incubator with 5% CO_2_ and routinely confirmed to be mycoplasma-free by PCR. All these cells were validated using short tandem repeat (STR) profiling.

### Lentivirus-mediated CAR construction

The fragments of GrB (21-247aa), CD40 (21-193aa), CD94 (31-179aa), NKG2A (94-233aa), NKG2C (94-231aa), TLR2 (21-588aa), and TLR4 (24-631aa) and the v-set domain of Siglec5 (101-136aa) were synthesized (BGI, China) and fused to a CAR backbone comprising the human CD8 hinge spacer and transmembrane domain, the 4-1BB costimulatory domain, and the CD3ζ signaling domain and fused to mKate2 via a T2A sequence. The entire coding sequence of the CAR expression molecule was cloned and inserted into the lentiviral vector pTomo (#26291, Addgene, USA) and expressed under the control of the CMV promoter. The CD19-specific scFv (FMC63) was inserted into the same backbone and serve as a negative control CAR.

For lentiviral packaging, the CAR-expressing plasmid was cotransfected with the packaging plasmid pCMVΔR 8.9 and envelope plasmid pMD2.G into 293T cells at a ratio of 10:5:2 using Lipo8000™ transfection reagent (Beyotime, China). The supernatant containing lentiviral particles was collected and concentrated by ultracentrifugation to collect lentivirus as previously described[45].

### CAR T-cell production

Human T cells were isolated from healthy donors who provided informed consent using RosetteSep Human T-Cell Enrichment Cocktail (STEMCELL Technologies, Canada) according to the manufacturer’s protocol. T cells were cultured in RPMI 1640 medium (Life Technologies) supplemented with 1×Glutamax (Life Technologies), 10% FBS, 1×penicillin-streptomycin (Life Technologies), and IL2 (200 U/ml, PeproTech) and stimulated with human CD3/CD28 Dynabeads (Life Technologies, USA) according to the manufacturer’s instructions. To generate CAR T cells, T cells were cultured for 72 h and infected with lentiviral particles in the presence of LentiBOOST (Sirion Biotech).

### *In vitro* cytotoxicity assays

The cytotoxicity of CAR T cells was evaluated using a luciferase assay system (Promega, USA) following the manufacturer’s protocol. In brief, NTD, CD19-CAR, or GrB-CAR T cells were cocultured with luciferase-labeled cancer cells for 16 h at different T-cell E:T ratios. The number of effector cells was calculated as the number of CAR-positive cells. Luciferase activity was measured, and cytotoxicity was calculated as the percentage of tumor cells treated with NTD T cells. To measure cytokine release upon cell lysis by CAR T-cell treatment, the supernatant was collected and analyzed with ELISA kits for IFNγ (Life Technologies) and TNFα (Proteintech, China) according to the manufacturer’s protocol.

### Quantitative PCR

Total RNA was isolated using TRI Reagent^®^ (Sigma-Aldrich) and treated with a URBO DNA-free Kit (Invitrogen) to remove contaminating genomic DNA according to the manufacturer’s instructions. RNA (2 μg) was reverse transcribed using random primers with a RevertAid First Strand cDNA Synthesis Kit (Thermo Scientific). Quantitative PCR was performed in triplicate using the SYBR Green method (Life Technologies). Relative expression levels were normalized to the level of 18S rRNA. To investigate the persistence of transduced T cells and tumor cells in blood, genomic DNA was purified from the blood by phenol/chloroform extraction for subsequent quantitative PCR with CAR-specific primers to determine the copy number of CAR T cells and with luciferase-specific primers to quantify tumor cells.

Human/mouse GAPDH were used as DNA loading controls as appropriate. All primers are listed in Table S1.

### Immunocytofluorescence

For mHSP70 staining, cancer cells (5×10^4^) were plated on coverslips in 24-well plates. The live cells were incubated with an anti-HSP70 antibody (StressMarq Biosciences, Canada) for 1 h, followed by washes and fixation with 4% PFA for 5 min. After that, the cells were washed three times and incubated with a goat anti-mouse secondary antibody (Life Technologies) for 1 h at room temperature in the dark. Nuclei were stained with 4’,6-diamidino-2-phenylindole (DAPI, 1 µg/ml, Sigma, USA) for 10 min in the dark.

### Flow cytometry

After being collected, the cells were resuspended in cold PBS with 2% FBS. The primary antibody was added to the cell suspension for incubation for 1 h. The cells were centrifuged, resuspended, and incubated with FITC-labeled secondary antibodies in PBS containing 2% FBS for 30 min prior to two washes with PBS. Then, flow cytometry was performed using a BD LSRFortessa cell analyzer (BD Biosciences, USA). The data were analyzed using FlowJo software. The antibodies used in the study are listed in Table S2.

### CAR T cell effects on xenograft growth and metastasis

The experimental protocols were approved by the animal ethics committee of the Kunming Institute of Zoology, Chinese Academy of Sciences. Six-week-old NOD/ShiLtJGpt-*Prkdc*^*em26Cd52*^*Il2rg*^*em26Cd22*^/Gpt (NCG) mice were purchased from GemPharmatech Co., Ltd. (Jiangsu, China). To explore the effects of GrB-CAR T cells on the tumor site, a total of 1×10^6^ AsPC-1-luc or SMMC7721-luc cells were suspended in PBS containing 30% Matrigel (Corning) and subcutaneously injected into NCG mice to establish the xenograft models. After 5 d, the mice were divided into two groups with similar mean bioluminescence signal intensities and treated with 3×10^6^ CD19-or GrB-CAR T cells.

In order to investigate the effects of GrB-CAR T cells on CTCs, NCG mice were subcutaneously injected with a total of 1×10^6^ luciferase-labeled AsPC-1 or SK-Hep-1 cells suspended in 200 μl of PBS containing 30% Matrigel. Based on bioluminescence measurement after 14 d, the xenografted mice were divided into 2 groups and treated with CD19-CAR T cells or GrB-CAR T cells by a single intravenous injection of gradient doses of 1×10^6^, 2×10^6^, and 3×10^6^ cells per mouse or by three intravenous injections of 5×10^5^ cells per mouse. Approximately 50 μl of blood was then collected from the animals at the indicated times to measure the copy number alterations in CAR T cells and tumor cells by quantitative PCR. The mice were sacrificed at the end of the experiment, and tissues were harvested to evaluate tumor metastasis in major organs, including the liver, lung, spleen, pancreas, kidney, and intestine. Tissues were fixed with 10% neutral buffered formalin and embedded in paraffin. Five-micrometer serial sections were stained with a human-specific anti-COXIV antibody and scanned using the Vectra® Polaris Automated Imaging System (PerkinElmer, USA) at 20×magnification. Images were analyzed in ImageJ, and metastases were counted.

### Enrichment and identification of CTCs

The spontaneous metastasis models was created utilizing AsPC-1 or SK-Hep-1 cells that stably expressed firefly luciferase and enhanced green fluorescent protein in order to detect CTCs in mice xenografts. A total of 1 ×10^6^ labelled cells were injected subcutaneously into NCG mice. Approximately 8 weeks after tumor cell inoculation, the mice were euthanized, and blood samples (approximately 300 μl) were collected via retroorbital bleeding. PBMCs were collected via RBC lysis and stained with an anti-HSP70 antibody and DAPI. The stained cells were seeded onto slides and subsequently subjected to scanning fluorescence microscopy.

Peripheral venous blood (6.0 ml) from each patient was drawn into a BD Vacutainer tube containing acid citrate dextrose (ACD) as an anticoagulant in order to detect CTCs in cancer patients (Becton Dickinson, Franklin Lakes, NJ, USA). The process for enrichment and identification of CTCs was performed with the Cytelligen CTC enrichment kit (Cytelligen, San Diego, CA, USA) following the manufacturer’s instructions. In brief, 6.0 ml peripheral venous blood was centrifuged to deplete serum, and the remaining components were mixed with 3 ml hCTC Separation Matrix. The mixture was centrifuged to remove RBCs, and the remaining cells were incubated with 150 μl immunomagnetic particles conjugated to anti-CD45 monoclonal antibodies for 20 min with gentle shaking at room temperature. CD45-positive leukocytes were removed using a magnetic stand. The remaining cells were washed 3 times with 1×CRC solution. Cell pellets were incubated with 100 µl of an anti-HSP70 antibody (1:100) for three hours at room temperature prior to washes and incubation with 200 µl of antibody preparation solution containing Alexa Fluor 594-conjugated anti-human CD45 for 20 min at room temperature in the dark. Next, the cells were seeded onto the slides and dried at 32 °C for 10 h for subsequent iFISH analysis with Centromere Probe 8 (CEP8) SpectrumOrange (Vysis, Abbott Laboratories, Abbott Park, IL, USA). The slides were stained with DAPI and subsequently subjected to scanning fluorescence microscopy. The study was approved by the Ethical Committee of West China Hospital, Sichuan University (Approval ID: 2022-858), and all patients signed informed consent forms.

### Immunohistochemistry

Sections were deparaffinized and rehydrated, and antigen retrieval was performed in citric acid solution (pH 6.0) for 15 min at 115 °C in a Decloaking Chamber^®^ (Biocare Medical). Following a 15-minute incubation in 3% hydrogen peroxide to inhibit endogenous peroxidase activity, the sections were blocked for 1 hour at room temperature using 10% goat serum. The sections were incubated overnight at 4 °C with primary antibodies, at room temperature for 1 h with fluorescence-labeled or HRP-labeled secondary antibodies, and with a Tyramide Signal Amplification (TSA) system (PerkinElmer). Nuclei were stained with DAPI prior to mounting with Aqua-Poly/Mount (Polysciences).

β-Catenin was used as a cell membrane marker to analyze the expression of mHSP70. Five images per tissue were acquired to quantify the colocalization of HSP70 and β-catenin by calculating Manders’ colocalization coefficients using ImageJ software with the Fiji plugin JACoP. Human tissue chips (HPanA060CS02 and HorgN120PT01) were purchased from Shanghai Outdo Biotech Co., Ltd. (China).

### Safety validation of CAR T cells in nonhuman primates

The rhesus macaques used in this study were obtained from the Kunming Primate Research Center at the Kunming Institute of Zoology, Chinese Academy of Sciences, and housed at the Association for Assessment and Accreditation of Laboratory Animal Care (AAALAC)-accredited facility of the Primate Research Center at Kunming Institute of Zoology. All protocols involving monkeys were approved by the animal ethics committee of the Kunming Institute of Zoology, Chinese Academy of Sciences (Approval ID: IACUC-PE-2021-05-002). PBMCs were isolated from three rhesus monkeys by density gradient centrifugation using Ficoll-Paque (STEMCELL Technologies). T cells were isolated from PMBCs using a Nonhuman Primate T-Cell Isolation Kit (STEMCELL Technologies), activated with a nonhuman primate T-Cell Activation/Expansion Kit (Miltenyi Biotec), and cultured in RPMI-1640 medium supplemented with 1×Glutamax (Life Technologies), 10% FBS, 1×penicillin-streptomycin sulfate (Life Technology) and IL-2 (200 U/ml, Pepro Tech). T cells were infected with GrB-CAR lentivirus in the presence of 1% LentiBOOST (Sirion Biotech GmbH), and 5 μg/ml cyclosporin A (Sigma-Aldrich). CAR T cells were intravenously injected into autologous monkeys (5×10^6^ total T cells/kg body weight in 1 ml saline). The monkeys were monitored daily for body temperature, weight, breathing rate, appetite, and diarrhea after injection of T cells. On d 0, 1, 3, 5, 7, 14, 21, 28, and 35, blood samples of approximately 1 ml were collected in EDTA-K2 anticoagulant tubes for analysis of hematological parameters using a BC-5000 Vet system. The blood samples were allowed to clot at room temperature in the absence of anticoagulants for serum harvesting. Serum biochemical parameters were measured using a Dimension® EXL™ 200 system. Using a macaque-specific multiplex assay (Meso Scale Discovery), Shanghai Universal Biotech Company determined the serum quantities of the cytokines TNF-α, IFN-γ, IL6, and IL2.

### Statistics

Statistical analysis was performed using GraphPad Prism software. Data are presented as the means ±SDs. Statistical comparisons were performed using Student’s *t*-test (two groups) or one-way analysis of variance (ANOVA) (three or more groups) with Tukey’s multiple comparison test. The levels of statistical significance were assigned as follows: not significant (ns), ^*^*P* ≤ 0.05, ^**^*P* ≤ 0.01, and ^***^*P* ≤ 0.001.

## Data availability

The main data supporting the results of this study are available within the paper and its Supplementary Information. Source data for the figures will be provided in this paper. All data generated in this study will be available from the corresponding author upon reasonable request.

## Acknowledgments

We want to thank Zhengfei Hu from the Kunming Primate Research Center, Kunming Institute of Zoology, Chinese Academy of Sciences for his technical support in nonhuman primate surgeries. We also gratefully acknowledge the technical assistance of the Core Facility of West China Hospital (Li Chai, Yi Li, Xing Xu, and Xiaoting Chen).

## Funding

This work was supported by grants from the National Natural Science Foundation of China (82172701 to X.Z., 92159302 to W. L.) and the 1.3.5 Project for Disciplines of Excellence, West China Hospital, Sichuan University (ZYYC20002 to X.Z.).

## Author contributions

X. Zhao, W. Li, and P. Shi designed the study, analyzed the data, wrote the manuscript, and were responsible for study supervision. B. Sun, J. Guo, H. Ma, N. Liu, and D. Hao designed the lentivirus vectors encoding CARs and performed *in vitro* experiments. B. Sun, D. Yang, and L. Lv performed monkey experiments. B. Sun, J. Guo, and L. Yan performed mouse experiments. D. Yang, Q. Hu, P. Tian, and N. Chen isolated and identified CTC from patient samples. H. Gou, D. Cao, and D. Liu provided technical and material support.

## Competing interests

X.Z. and B.S are inventors in a patent filed by the West China Hospital, Sichuan University (application number: 202110667439.X). The other authors declare that they have no competing interests.

