## Supplementary Figures 1-15, Table S1, Table S2 for "Granzyme B-based CAR T cells block metastasis by eliminating circulating tumor cells"

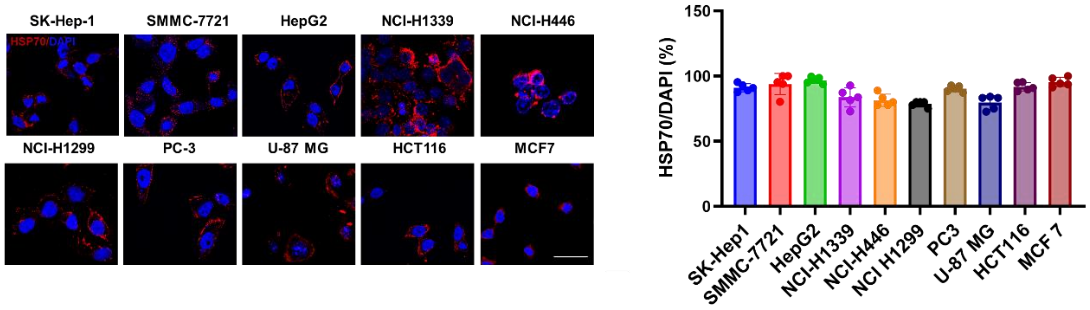


**Supplementary Fig. 1 Expression of mHSP70 in various cancer cell lines**

Representative images of mHSP70 expression in SK-Hep1,SMMC-7721,HepG2, NCI-H1339, NCI-H446, NCI-H1299, PC3, U-87 MG, HCT116, and MCF7 cells, as detected by immunofluorescence (left). Scale bars, 50 μm. The percentage of mHSP70-positive cells was quantified (right), n=5 independent experiments.

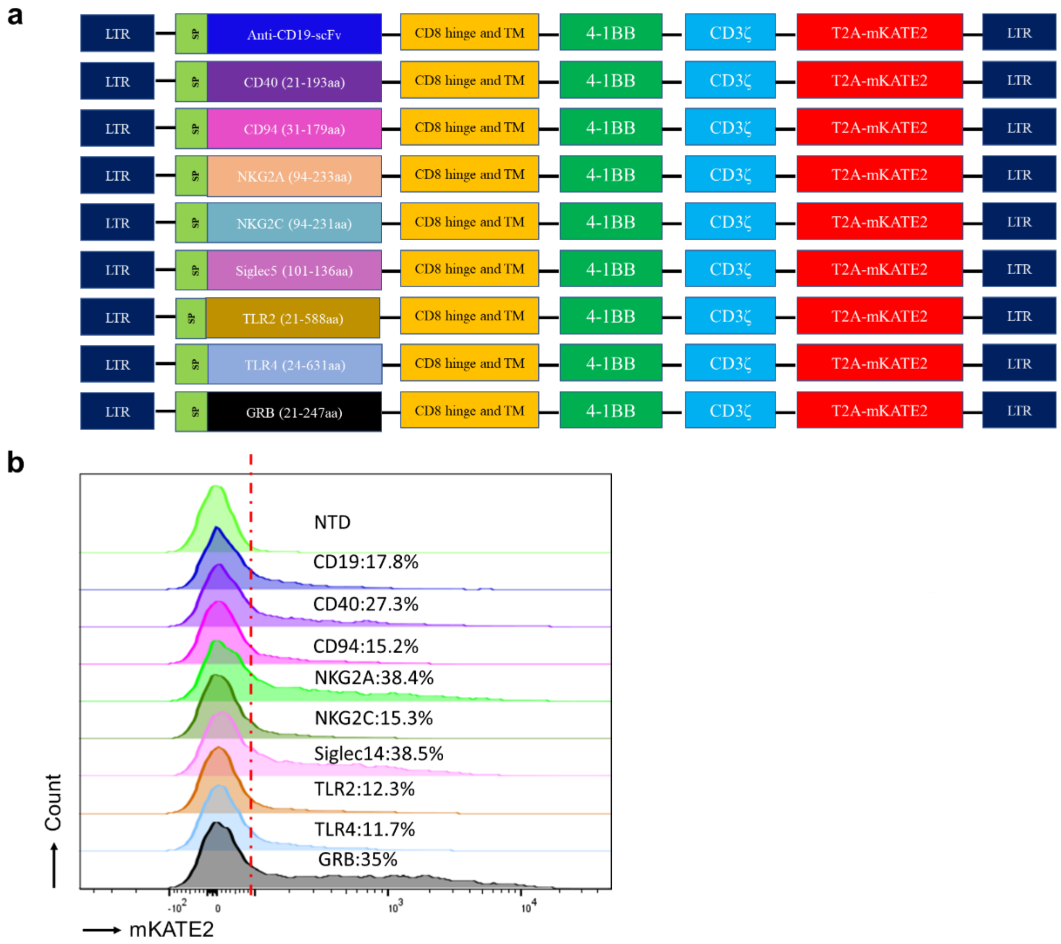


**Supplementary Fig. 2 Optimization of the GrB-CAR design for effector potency**

**a**, Schematic representation of the CAR constructs targeting mHSP70. **b,** Flow cytometric analysis of CAR expression compared to control (NTD).

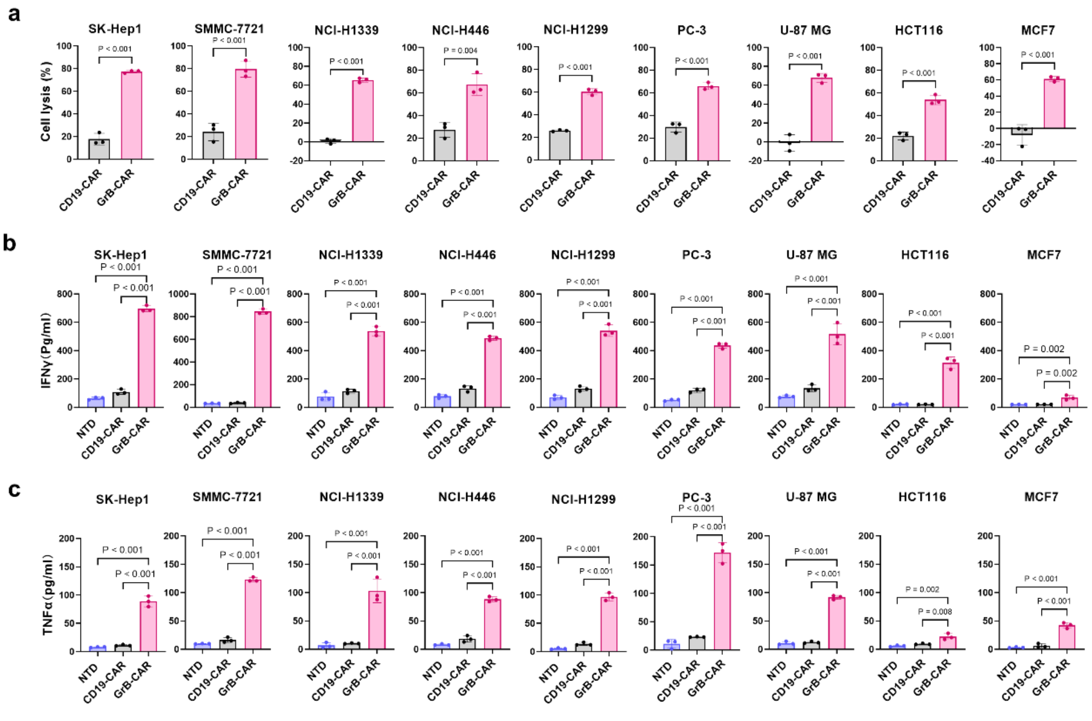


**Supplementary Fig. 3 Cytotoxic effect of GrB-CAR T cells in multiple cancer cell lines**

**a,** Cytotoxicity of GrB-CAR T cells against the indicated cells at an E/T ratio of 3:1. **b, c**,Release of the cytokines IFN-γ (**b**) and TNF-α (**c**) as determined by ELISA in the supernatant of various cancer cells treated with control or CAR T cells. The data are presented as the means ± SDs. Data in a-c are representative of three biological replicates from n=2 independent experiments.

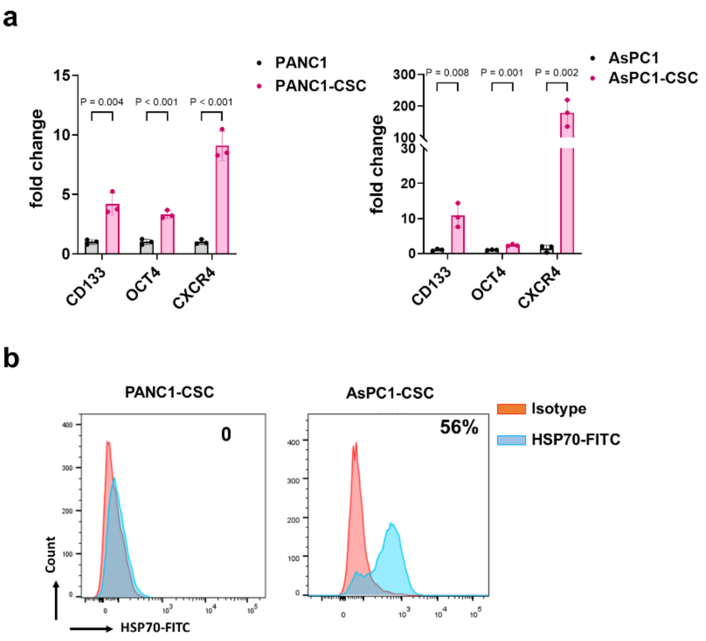


**Supplementary Fig. 4 GrB-CAR T cells target stem-like cancer cells expressing mHSP70**

**a,** Expression of the cancer stem cell markers CD133, OCT4, and CXCR4 in AsPC1-CSCs and PANC1-CSCs as determined by quantitative real-time PCR. The data are presented as the means ± SDs. Data are representative of three biological replicates from n=2 independent experiments. **b,** Flow cytometric analysis of mHSP70 expression on stem-like AsPC1-CSCs and PANC1-CSCs. Data are representative of two independent experiments.

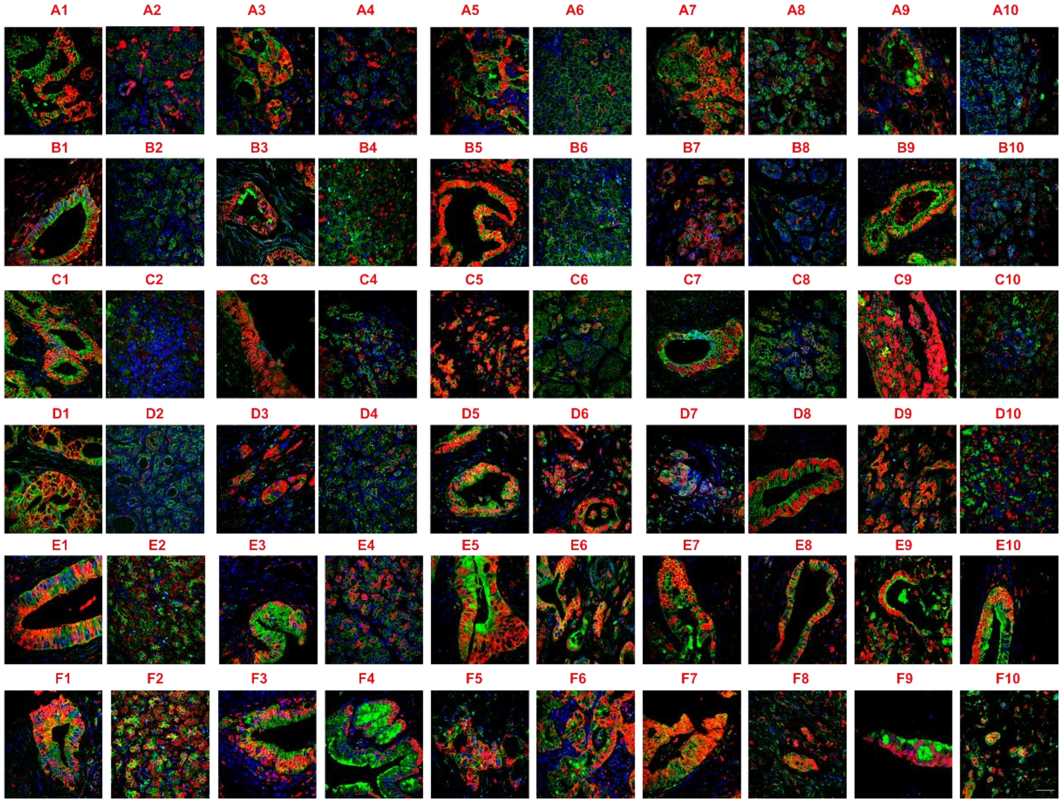


**Supplementary Fig. 5 The expression of mHSP70 in human pancreatic tumor tissues**
Representative images of costaining of mHSP70, β-catenin and DAPI in a tissue microarray containing 23 pairs of pancreatic cancer and paracancerous tissue (A1 to E6; odd numbers, cancer tissue; even numbers, paracancerous tissue), as well as 14 additional pancreatic tissue samples (E7 to F10). HSP70, red; β-catenin, green; nuclei-DAPI, blue. Scale bars, 40 μm.

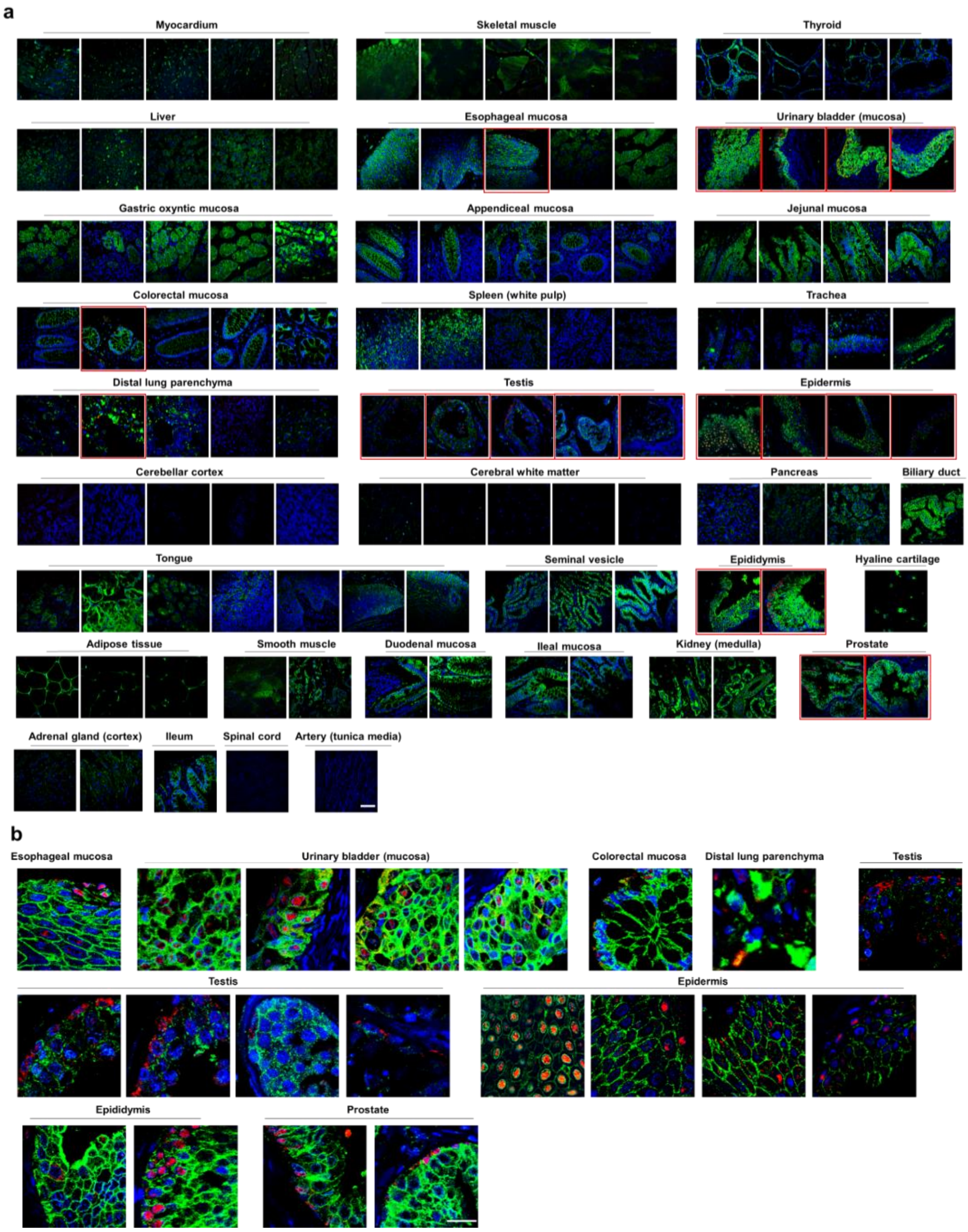


**Supplementary Fig. 6 The expression of mHSP70 in normal human tissues**

**a, b**, Representative images of costaining of mHSP70, β-catenin and DAPI in a tissue microarray containing 37 normal human samples. The tissues expressing HSP70 are identified by red boxes, and the images are enlarged and displayed in (b). HSP70, red; β-catenin, green; nuclei-DAPI, blue. Scale bars in (a), 50 μm; scale bars in (b), 20 μm.

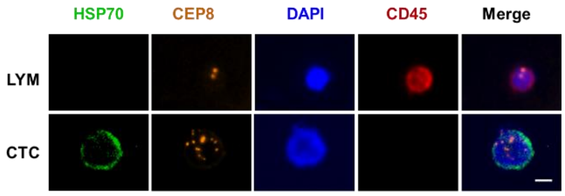


**Supplementary Fig. 7 Expression of mHSP70 on CTCs derived from human pancreatic tumors.**

Representative images of mHSP70 expression on CTCs derived from pancreatic cancer. HSP70, green; CEP8, orange; CD45, red, and nuclei-DAPI, blue. Scale bars, 10 μm.

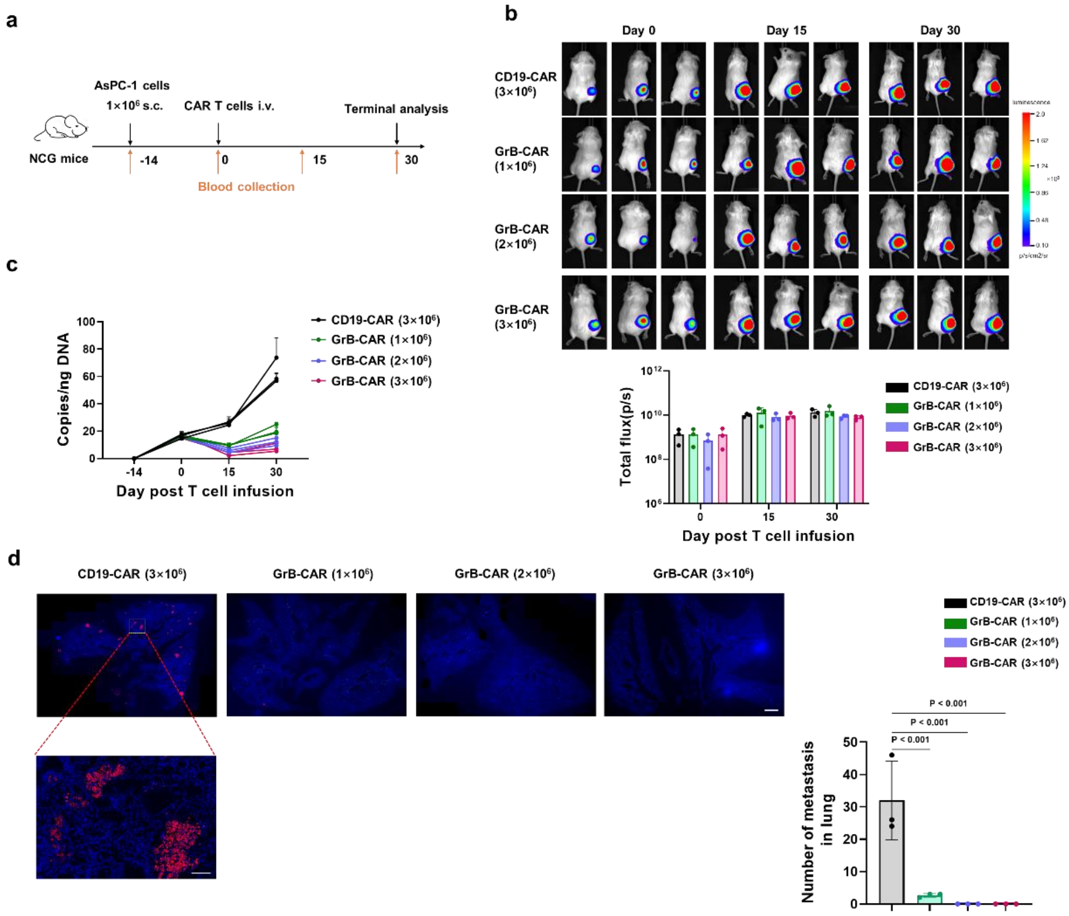


**Supplementary Fig. 8 A low number of GrB-CAR T cells suppresses tumor metastasis in an AsPC1 xenograft model.**

**a,** Schematic diagram of the in vivo experimental model. **b,** In vivo bioluminescence imaging of tumor growth in mice treated with CD19 and GrB-CAR T cells at the indicated times (upper); statistical analysis of tumor bioluminescence (lower). **c,** Detection of DNA copies of CTCs in peripheral blood collected from tumor-bearing mice. n=3. **d,** Metastasis of human tumor cells in mouse lung tissue upon CAR T-cell challenge as indicated. Representative images of immunofluorescence staining with a human-specific anti-COXIV antibody (red) and DAPI (blue) in the lung (left). Upper scale bars, 800 μm; lower scale bars, 100 μm. Statistical analysis of the tumor nodules (right). Each dot represents the metastatic nodes in an individual mouse. The data are shown as the means ± SDs, n=3.

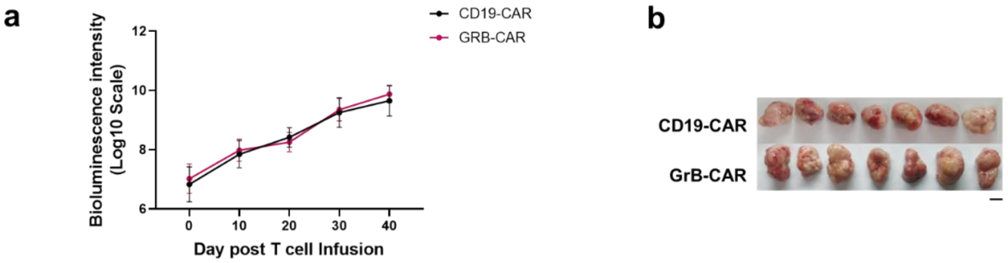


**Supplementary Fig. 9 GrB-CAR T cells suppress tumor metastasis without affecting the growth of AsPC-1 xenograft tumors in NCG mice.**

**a,** NCG mice bearing AsPC1-luc cells were injected with CD19-CAR T or GrB-CAR T cells. Quantification of tumor bioluminescence levels at different time points. n=7; ns, not significant. **b,** Images of the tumors in NCG mice 40 days after CAR T-cell treatment.

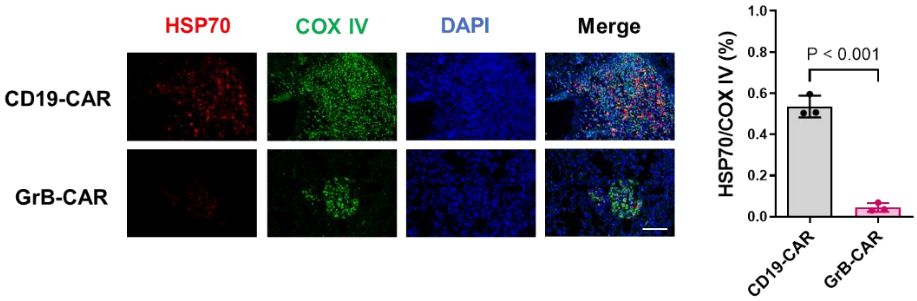


**Supplementary Fig. 10 The expression of HSP70 in AsPC-1 metastatic tumors**

Left, representative images of immunofluorescence staining with HSP70 (red), a human-specific anti-COXIV antibody (green), and DAPI (blue) in lungs harvested from mice on day 40 after treatment with CD19-CAR or GrB-CAR T cells. Scale bar, 100 μm. The percent of $HSP70^+$ cells among $COXIV^+$ cells in tumor tissues was quantified based on five images per sample, n=3 per group (right). The data are shown as the means ± SDs.

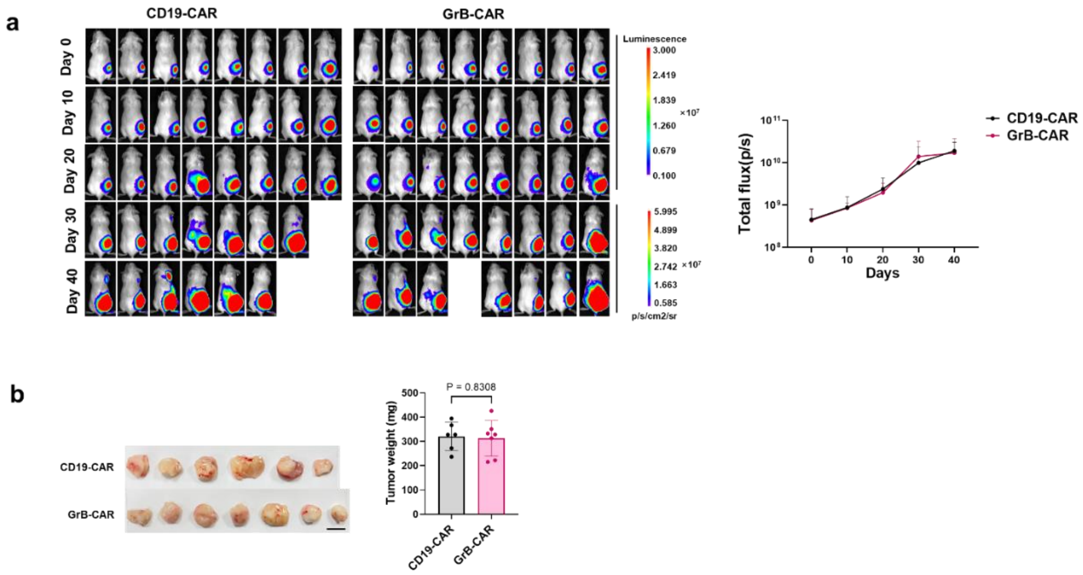


**Supplementary Fig. 11 GrB-CAR T cells suppress tumor metastasis without affecting the growth of SK-Hep1 xenograft tumors in NCG mice.**

**a,** Left, bioluminescence imaging of mice bearing tumors derived from the human hepatoma cell line SK-Hep1 and treated with CD19-CAR or GrB-CAR T cells. Each column shows one mouse over time. Right, quantification of tumor bioluminescence at different time points (right). **b,** Tumor weight in NCG mice was quantified 40 days after CAR T-cell treatment. The data are presented as the means ± SDs. CD19-CAR, n=6; GrB-CAR, n=7.

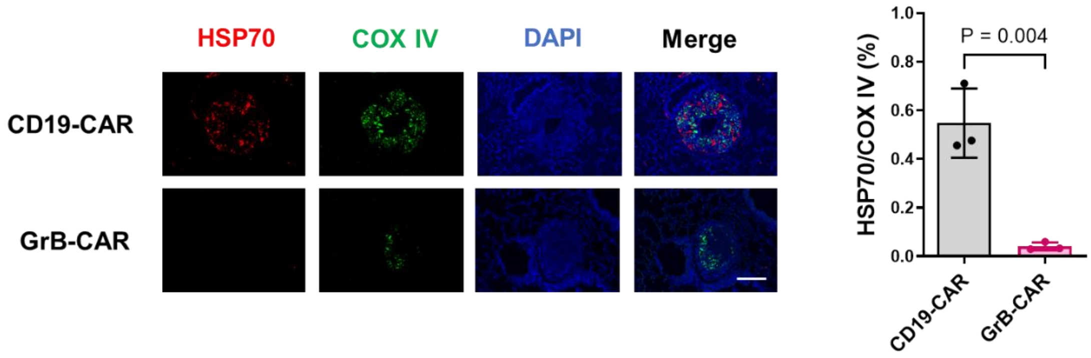


**Supplementary Fig. 12 The expression of HSP70 in SK-Hep1 metastatic tumors**

Left, representative images of immunofluorescence staining with HSP70 (red), a human-specific anti-COXIV antibody (green), and DAPI (blue) in lungs harvested from mice on day 40 after treatment with CD19-CAR or GrB-CAR T cells. Scale bar, 100 μm. The percent of $HSP70^+$ cells among $COXIV^+$ cells in tumor tissues was quantified based on five images per sample, n=3 per group (right). The data are shown as the means ± SDs.

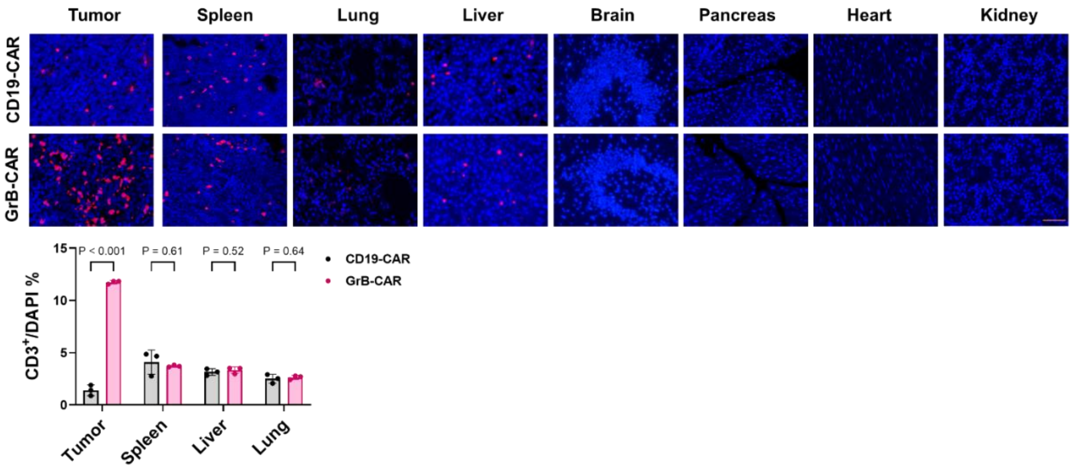


**Supplementary Fig. 13 CAR T cell infiltration in tissues from mice treated with CAR T cells**

Representative images of CAR T cells in various organs are shown. NCG mice bearing AsPC-1 cells were injected with CD19 CAR T cells or GrB-CAR T cells. After 5 days, the mice were sacrificed, and the indicated organs were collected for CD3 staining (*red*, upper). Scale bars, 100 μm. Quantification of $CD3^+$ T cells per image field is shown as the mean ±SD (lower), $n$=3 for each group.

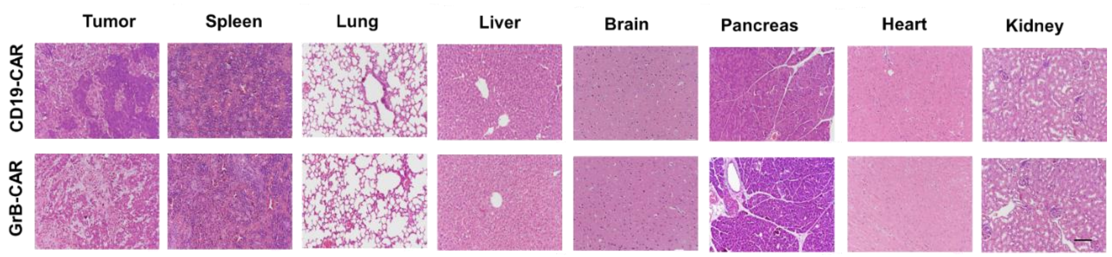


**Supplementary Fig. 14 Pathological changes in tissues from mice treated with CAR T cells**

Representative histological images of various organs are shown. NCG mice bearing AsPC-1 cells were injected with CD19 CAR T cells or GrB-CAR T cells. After 5 days, the mice were sacrificed, and the indicated organs were collected for H&E staining (*n*=3 mice per group). Scale bars, 100 μm.

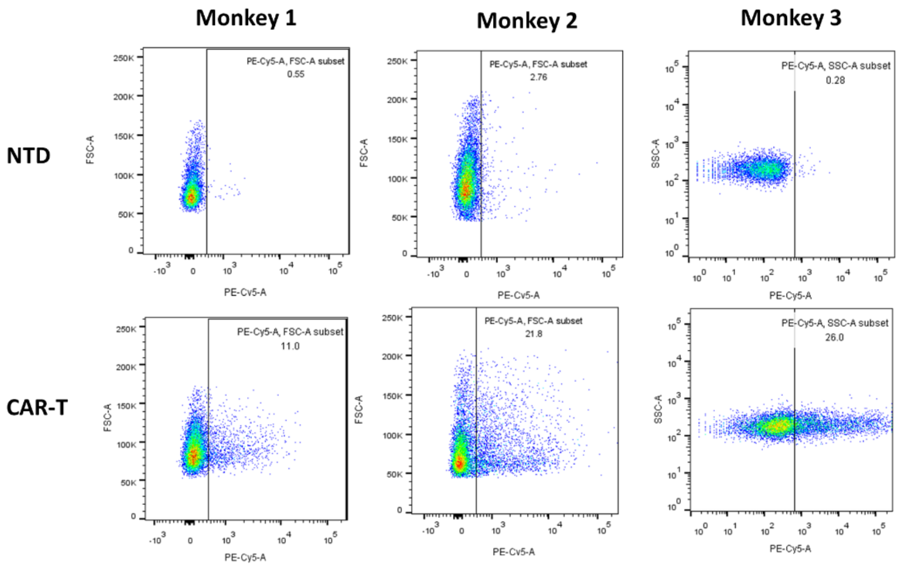


**Supplementary Fig. 15 CAR efficiency of nonhuman primate GrB-CAR T cells.** GrB-CAR expression in T cells of nonhuman primates was evaluated by flow cytometry 72 h after transduction. The percentages represent the numbers of CAR-positive cells. Nontransduced T cells were used as controls. The data are representative of n=2 independent experiments.

### Table S1. List of Realtime PCR primers

| Name | Organism | Sequence |
|---|---|---|
| RPS18-F | Human | GTAACCCGTTGAACCCCATT |
| RPS18-R | Human | CCATCCAATCGGTAGTAGCG |
| HSPA1A-F | Human | ACCTTCGACGTGTCCATCCTGA |
| HSPA1A-R | Human | TCCTCCACGAAGTGGTTCACCA |
| CD133-F | Human | AGTCGGAAACTGGCAGATAGC |
| CD133-R | Human | GGTAGTGTTGTACTGGGCCAAT |
| CXCR4-F | Human | AAACTGAGAAGCATGACGGACAA |
| CXCR4-R | Human | GCCAACATAGACCACCTTTTCAG |
| OCT4-F | Human | GTGCCGTGAAGCTGGAGAA |
| OCT4-R | Human | TGGTCGTTTGGCTGAATACCTT |
| Gapdh-F | Mouse /Human | CATCACTGCCACCCAGAAGACTG |
| Gapdh-R | Mouse /Human | ATGCCAGTGAGCTTCCCGTTCAG |
| GAPDH-F | Monkey | CTGGGCTACACTGAGCACC |
| GAPDH-R | Monkey | AAGTGGTCGTTGAGGGCAATG |
| GRB-CD8-F | Human | CCAGGGCATTGTCTCCTATGG |
| GRB-CD8-R | Human | TCACAGGCGAAGTCCAGC |
| Luc-F | | TGCGCGGAGGAGTTGTGTTTGTG |
| Luc-R | | ACGGCGATCTTTCCGCCCTTCT |
| U5-F | | AGCTTGCCTTGAGTGCTTCA |
| U5-R | | TGACTAAAAGGGTCTGAGGG |

### Table S2. Summary of antibody

| Name | Company | Catalog | Species | Usage | Dilution ratio |
|---|---|---|---|---|---|
| HSP70 | Stressmarq | SMC-249 | Human | WB | 1:1000 |
| | | | | IHC | 1:100 |
| | | | | IF | 1:100 |
| | | | | FCM | 1:250 |
| β-catenin | HUABIO | ER0805 | Human | IHC | 1:200 |
| CD3-episilon | HUABIO | ET1607-29 | Human | IHC | 1:200 |
| COXIV | Cell Signaling Technology | #4850 | human | IHC | 1:1000 |
